# IL-11 neutralising therapies target hepatic stellate cell-induced liver inflammation and fibrosis in NASH

**DOI:** 10.1101/470062

**Authors:** Anissa A. Widjaja, Brijesh K. Singh, Eleonora Adami, Sivakumar Viswanathan, Giuseppe A. D’Agostino, Jinrui Dong, Benjamin Ng, Jessie Tan, Bhairav S. Paleja, Madhulika Tripathi, Sze Yun Lim, Sonia P. Chothani, Wei Wen Lim, Anne Rabes, Martina Sombetzki, Eveline Bruinstroop, Rohit A. Sinha, Salvatore Albani, Paul M. Yen, Sebastian Schafer, Stuart A. Cook

**Author notes:** Correspondence: S.A.C. These authors contributed equally to this work (Anissa A. Widjaja, Brijesh K. Singh).

## Abstract

The transformation of hepatic stellate cells (HSCs) into myofibroblasts is the defining pathobiology in non-alcoholic steatohepatitis (NASH). Here we show that key NASH factors induce IL-11, which drives an autocrine and ERK-dependent activation loop to initiate and maintain HSC-to-myofibroblast transformation, causing liver fibrosis. IL-11 is upregulated in NASH and *Il11ra1*-deleted mice are strongly protected from liver fibrosis, inflammation and steatosis in murine NASH. Therapeutic inhibition of IL11RA or IL-11 with novel neutralizing antibodies robustly inhibits NASH pathology in preclinical models and reverses established liver fibrosis by promoting HSC senescence and favourable matrix remodelling. When given early in NASH, IL-11 inhibition prevents liver inflammation and steatosis, reverses severe hepatocyte damage and reduces hepatic immune cells and TGFβ1 levels. Our findings show an unappreciated and central role for IL-11 in HSCs and prioritise IL-11 signalling as a new therapeutic target in NASH while revealing an unexpected pro-inflammatory function for IL-11 in stromal immunity.

## Introduction

The global prevalence of nonalcoholic fatty liver disease (NAFLD) is estimated at 25%^1^ and while NAFLD is reversible it can progress to nonalcoholic steatohepatitis (NASH). NASH is characterized by steatosis-driven inflammation, hepatocyte death and liver fibrosis that can lead to liver failure. Hepatic stellate cells (HSCs) are pivotal in the pathogenesis of NASH and give rise to up to 95% of liver myofibroblasts^2^, which drive the key pathologies in NASH, namely liver fibrosis, inflammation and dysfunction^3–5^.

A number of factors are implicated in HSC activation and transformation, including the canonical pro-fibrotic factors transforming growth factor‑β1 (TGFβ1) and platelet-derived growth factor (PDGF)^6,7^ and also pro-inflammatory factors such as CCL2, TNFα and CCL5^4,7,8^. Perhaps reflecting this redundancy in HSC activation, no single upstream initiating factor has been targeted successfully in NASH. Inhibition of downstream pro-fibrotic targets such as LOXL2 has also been unsuccessful and ongoing clinical trials are focused mostly on inhibiting steatosis. There are no approved drugs for the treatment of NASH.

Quiescent HSCs are vitamin A storing cells and very distinct from fibroblasts. However, common stimuli activate both cell types and stimulate their transition to myofibroblasts with shared features^2,9^. We recently identified Interleukin-11 (IL-11) as a crucial factor for cardiovascular and pulmonary fibroblast-to-myofibroblast transformation^10,11^. To date, there are very limited insights into IL-11 in the liver, where it is reported to have anti-inflammatory activity^12,13^, and it is unknown if HSCs respond to IL-11 at all. Here, we explore the hypothesis that IL-11 plays a role in the transformation of HSCs into myofibroblasts and determine the effects IL-11 signalling in the context of liver inflammation, steatosis and fibrosis in NASH.

### IL-11 activates HSCs and drives liver fibrosis in NASH

Genome wide RNA-seq analysis revealed that TGFβ1 strongly upregulates *IL-11* (14.9-fold, P = 3.40×10^−145^) in HSCs that was confirmed by qPCR and at the protein level and replicated in experiments using precision cut human liver slices (**Fig. 1a-c, Supplementary Fig. 1a**). Independent RNA-seq data^14^ also show that *IL-11* is the most upregulated gene in HSCs when grown on a stiff substrate to model cirrhotic liver (**Supplementary Fig. 1b**). HSCs express very low levels of IL6R and higher levels of the IL-11 receptor subunit alpha (IL11RA) than either cardiac or lung fibroblasts (**Fig. 1d, Supplementary Fig. 1c**). We also performed Western blots on patient liver samples and found increased IL-11 levels in NASH (**Supplementary Fig. 1d,e**). These data show that HSCs are both a source and prominent target of IL-11 in the human liver and that IL-11 is elevated in NASH.

**Figure 1.**
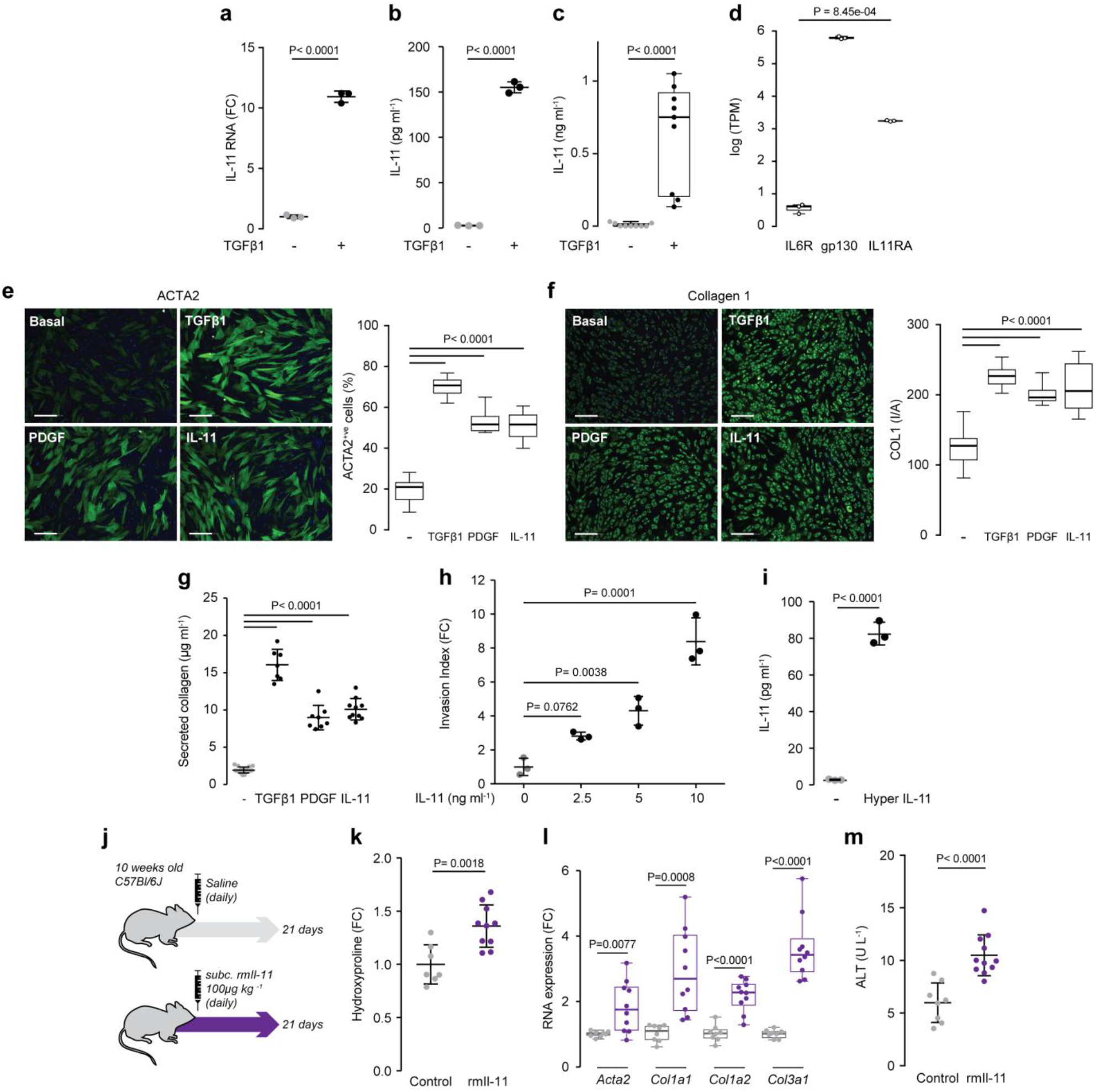
IL-11 induces hepatic stellate cell activation and hepatic fibrosis. **a**, *IL-11* is upregulated in hepatic stellate cells (HSCs) stimulated with TGFβ1 (n=3). **b**, IL-11 protein is secreted from HSCs stimulated with TGFβ1 (ELISA, n=3). **c**, Human precision cut liver slices were stimulated with TGFβ1 and IL-11 protein was measured in supernatant (ELISA, n=3). **d**, IL6R, gp130, and IL11RA expression in HSCs (TPM, transcripts per million). **e, f**, Representative fluorescence images (scale bars, 200 µm) of HSCs and automated fluorescence quantification for (**e**) ACTA2^+ve^ cells and (**f**) Collagen I immunostaining following incubation without stimulus (-), with TGFβ1, PDGF, or IL-11. **g**, Collagen secretion supernatants of HSC stimulated with TGFβ1, PDGF, or IL-11 (Sirius red assay. n≥7). **h**, Dose-dependent matrigel invasion of HSCs induced by IL-11(n=3). **i**, Hyper IL-11 induces IL-11 protein secretion from HSCs (ELISA, n=3). **a-c, e-g, i**, TGFβ1 (5 ng ml^-1^), Hyper IL-11 (0.2 ng ml^-1^), PDGF (20 ng ml^-1^), IL-11 (5 ng ml^-1^); 24 h stimulation; **h**, 48 h stimulation. **j**, Schematic of mice receiving daily subcutaneous injection of either saline (control) or rmIl-11 (100 µg kg^-1^). **k**, Relative liver hydroxyproline content, **l**, mRNA expression of pro-fibrotic markers, and **m**, serum ALT levels **(k, l**, control, n=7; rmIl-11, n=10; **m**, control n=8; rmIl-11, n=11). **a, b, g, h, i, k, m** Data are represented as mean ±s.d; **c-f, l**, Box-and-whisker plots show median (middle line), 25th–75th percentiles (box) and min-max percentiles (whiskers). **a-d, i, k-m**, Two-tailed Student’s *t*-test; **e-h**, two-tailed Dunnett’s test. FC: fold change; I/A: intensity/area.

To investigate the effect of IL-11 on HSCs, we stimulated cells with either IL-11, TGFβ1 or PDGF. IL-11 activated HSCs to a similar extent as TGFβ1 or PDGF, transforming quiescent HSCs into ACTA2^+ve^ myofibroblasts that secrete collagen and matrix modifying enzymes (**Fig. 1e-g, Supplementary Fig. 1f**). IL-11 also promoted dose-dependent matrix invasion by HSCs that is an important aspect of HSC pathobiology in NASH (**Fig. 1h**). We stimulated HSCs with with hyperIL-11^10^ to test for an autocrine loop of feed-forward IL-11 activity, inferred by IL-11 secretion from HSCs that express IL11RA, and confirmed its existence (**Fig. 1i**). Moving *in vivo*, subcutaneous administration of recombinant mouse Il-11 (rmIl-11) to mice for 21 days increased hepatic collagen content, fibrosis marker mRNA and serum alanine aminotransferase (ALT) levels (**Fig. 1j-m**). Furthermore, *Col1a1-GFP* reporter mice^15^ treated with rmIl-11 accumulated *Col1a1*^+ve^ myofibroblasts in the liver (**Supplementary Fig. 1g**).

### Anti-IL-11 therapies are effective in treating murine NASH

We next performed studies in a murine model of severe NASH using the high fat methionine- and choline-deficient (HFMCD) diet^16^. In this model, *Il-11* mRNA was mildly elevated whereas protein levels were highly upregulated, revealing strong post-transcriptional regulation of Il-11 expression in the liver (**Supplementary Fig. 2a,b**). The progressive induction of Il-11 protein was mirrored by ERK activation, increased collagen and elevated serum ALT levels (**Fig. 2a-c, Supplementary Fig. 2c**). To evaluate the physiological relevance of increased Il-11 levels in NASH, we used a genetic loss-of-function model: the Il-11 receptor subunit alpha deleted mouse (*Il11ra1^-/-^*)^17^. *Il11ra1^-/-^* mice on the NASH diet were protected from fibrosis and had lesser steatosis and liver damage and ERK activation (**Fig. 2d-g, Supplementary Fig. 2d,e**). Hence, non-canonical and ERK-dependent Il-11 signalling, seen previously during fibroblasts-to-myofibroblast transformation^10^, appeared relevant for several distinct aspects of NASH pathobiology.

**Figure 2.**
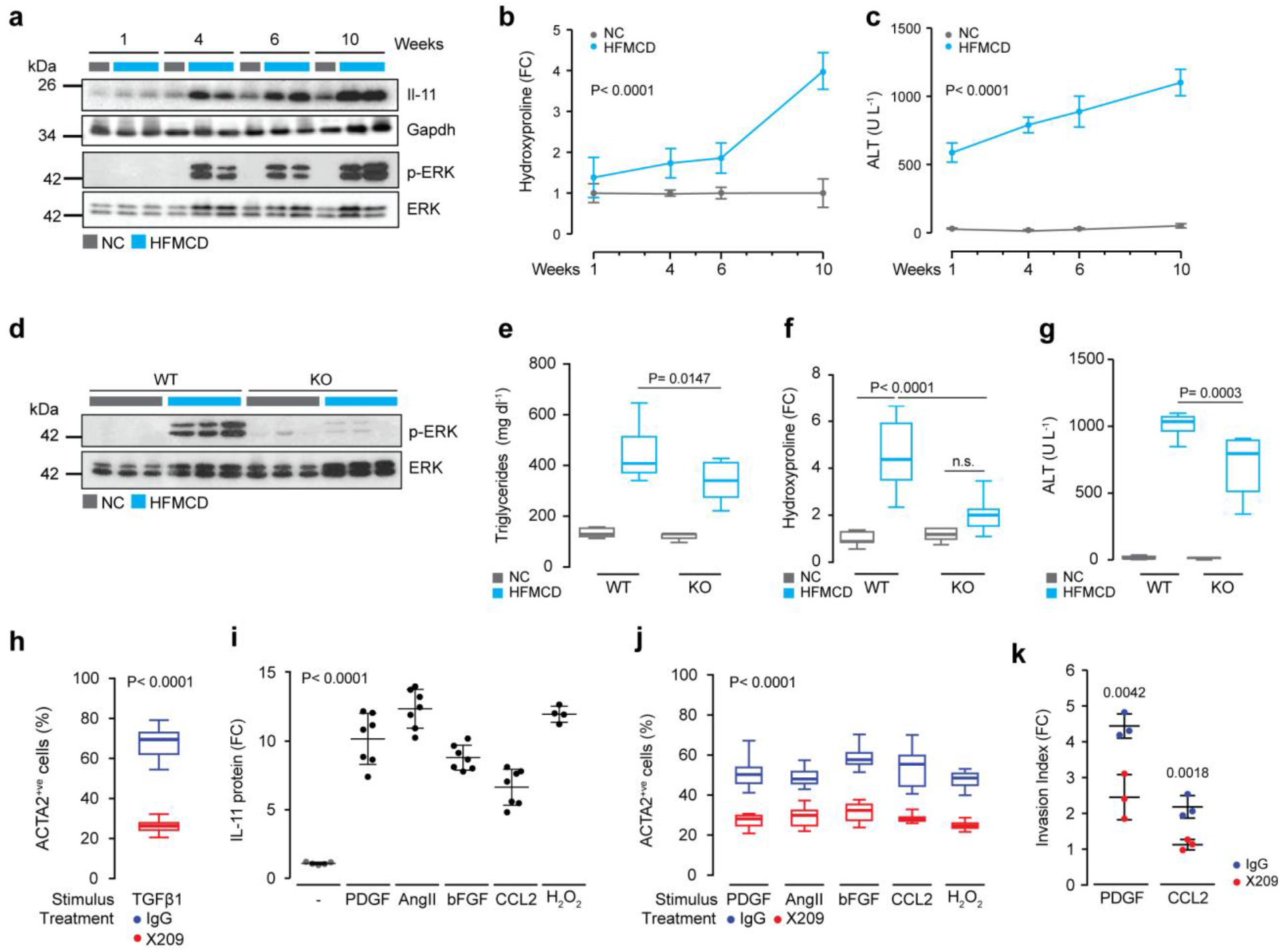
Inhibition of Il-11 signalling prevents hepatic stellate cell activation and hepatic fibrosis. **a**, Western blots of hepatic Il-11 and ERK activation status in mice on HFMCD diet. **b**, Relative liver hydroxyproline content and **c**, serum ALT levels in mice fed on HFMCD diet; liver tissues and serum were collected at the indicated time points (NC 1, 4, 6 week(s), n=5; NC HPA 10 weeks, n=4; NC ALT 10 weeks, n=5; HFMCD 1 week, n=7; HFMCD 4, 6 weeks, n=8; HFMCD 10 weeks, n=5). **d**, Western blots of hepatic ERK activation status after 10 weeks of HFMCD diet in *Il11ra^+/+^* (WT) and *Il11ra^-/-^* (KO) mice. **e**, Liver triglyceride levels and **f**, relative liver hydroxyproline content in WT and KO mice fed with HFMCD diet for 10 weeks (NC WT, n=9; HFMCD WT, n=9; NC KO, n=5; HFMCD KO, n=9). **g**, Serum ALT levels in WT and KO mice following HFMCD diet (NC WT, n=9; HFMCD WT, n=8; NC KO, n=5; HFMCD KO, n=9). **h**, ACTA2^+ve^ cells numbers in hepatic stellate cell (HSC) cultures stimulated with TGFβ1 in the presence of either IgG or X209. **i**, ELISA of IL-11 secretion from HSCs stimulated with various NASH factors (basal, n=5; PDGF, AngII, bFGF, CCL2, n=7; H_2_O_2_, n=4). **j**, Effect of X209 on ACTA2^+ve^ cell proportions in HSCs stimulated with various NASH factors. **k**, Effects of X209 on PDGF- or CCL2-induced HSC invasion (n=3). **h, j**, 24 h stimulation; **i, k**, 48 h stimulation. **h-k**, TGFβ1 (5 ng ml^-1^), Hyper IL-11 (0.2 ng ml^-1^), PDGF (20 ng ml^-1^), AngII (100 nM), bFGF (10 ng ml−1), CCL2 (5 ng ml^-1^), H 0 (0.2 mM), IgG and X209 (2 μg ml^-1^). **b, c, i, k**, Data are represented as mean ±s.d; **e-h, j**, data are shown as box-and-whisker with median (middle line), 25th–75th percentiles (box) and min-max percentiles (whiskers). **b, c**, Two-way ANOVA; **e-g**, two-tailed, Sidak-corrected Student’s *t*-test; **h-k**, two-tailed Dunnett’s test. FC: fold change; NC: normal chow; HFMCD: high fat methionine- and choline-deficient.

In an attempt to target the IL-11 autocrine loop, we genetically immunised mice with IL11RA to generate neutralising anti-IL11RA antibodies. Clones that block fibroblast transformation^10^ were identified and clone X209 (IgG1_K_, *K*_D_ = 6nM) that neutralised IL-11 signalling across species was prioritised. X209 blocked fibrogenic protein secretion from HSCs with an IC_50_ of 5.4pM. Pharmacokinetic studies using ^125^I-X209 revealed an *in vivo* half-life of more than 18 days and strong uptake in the liver (**Supplementary Fig. 3**). To ensure therapeutic specificity for IL-11 signalling and exclude off-target effects, we also developed a neutralising anti-IL-11 antibody (X203)^11^ and used both antibodies in downstream studies.

Initial experiments revealed that both antibodies blocked the TGFβ1-driven transition of HSCs into myofibroblasts (**Fig. 2h, Supplementary Fig. 4a,b,d,e,g**). Follow on studies found that other key NASH stimuli such as PDGF, CCL2, angiotensin II, bFGF or oxidative stress also induce IL-11 secretion from HSCs. And, remarkably, HSC-to-myofibroblast transformation downstream of these various stimuli is consistently dependent on intact IL-11 signalling (**Fig. 2i-k, Supplementary Fig. 4c,f,h,i**). Thus, IL-11 activity is a critical and universal feature underlying HSC transformation, which has similarities with its activity in fibroblasts^10,11^.

We then tested X209 and X203 therapy *in vivo* and started antibody administration (10 mg kg^-1^ bi-weekly) after six weeks of NASH diet when IL-11 is strongly upregulated, collagen has accumulated and there is severe steatohepatitis (**Fig. 2a-c, 3a, Supplementary Fig. 5a**). After four weeks of therapy both antibodies had abolished ERK activation, demonstrating excellent target engagement and coverage. Anti-IL-11 therapies inhibited the progression in liver fibrosis and serum ALT levels, while steatosis was largely unchanged (**Fig. 3b-e**, **Supplementary Fig. 5b-g)**.

**Figure 3.**
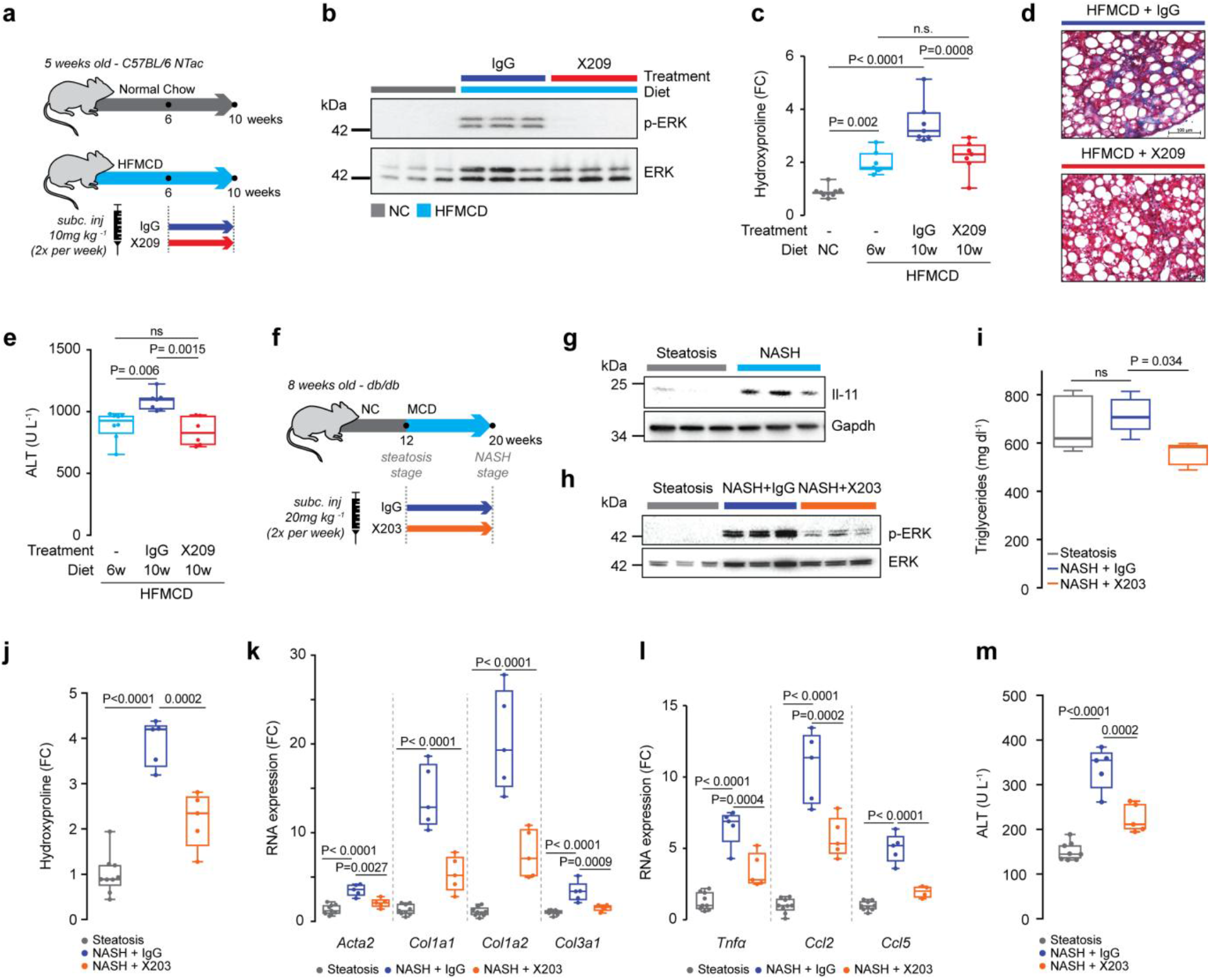
Therapeutic inhibition of Il-11 signalling inhibits the progression of late stage NASH. **a**, Schematic showing therapeutic use of X209 in HFMCD-fed mice. X209 or IgG isotype control (10 mg kg^-1^, twice a week) were administered from week 6 to 10 of HFMCD diet. **b-e**, Data for therapeutic dosing experiments as shown in **3a. b**, Western blots of hepatic ERK activation status. **c**, Relative liver hydroxyproline content (NC, n=9; HFMCD 6 weeks, n=8; IgG, n=8; X209, n=9; the values of NC and HFMCD 6 weeks are the same as those used in **2b**), **d**, representative Masson’s Trichrome staining, **e**, serum ALT levels (HFMCD 6 weeks, n=8; IgG, n=7; X209, n=6; the values of HFMCD 6 weeks are the same as those used in **2c**) of X209- and IgG-treated mice. **f**. Schematic of X203 or IgG administration to MCD-fed *db*/*db* mice and times of liver sample collection for use in experiments shown in **g**-**m**. **g**, Western blots of hepatic Il-11 and Gapdh. **h**, Western blots of total and phosphorylated ERK levels in livers of X203 or IgG-treated mice. **i**, Hepatic triglyceride content, **j**, liver hydroxyproline content, and **k**, pro-fibrotic and **l**, pro-inflammatory mRNA expression (steatosis, n=9; NASH+IgG, n=5; NASH+X203, n=5). **m**, Serum ALT levels (steatosis, n=8; NASH+IgG, n=5; NASH+X203, n=5). **c, e, i-m**, Data are shown as box-and-whisker with median (middle line), 25th– 75th percentiles (box) and min-max percentiles (whiskers); two-tailed, Tukey-corrected Student’s *t*-test. FC: fold change; NC: normal chow; HFMCD: high fat methionine- and choline-deficient; MCD: methionine- and choline-deficient.

To extend these findings, we tested anti-IL-11 therapy in an additional NASH model using obese and insulin resistant (*db/db*) mice on a methionine- and choline-deficient (MCD) diet (**Fig. 3f**)^18–20^. As expected, Il-11 expression and ERK activation were increased in livers of MCD fed *db/db* mice (**Fig. 3g,h**). Furthermore, anti-IL11 therapy reduced hepatic steatosis, fibrosis, inflammation and improved ALT levels as compared to controls (**Fig. 3i-m, Supplementary Fig. 6a**). A third model of streptozotocin-induced diabetes and advanced NASH (**Supplementary Fig. 6b**) was investigated although ALT was not elevated in this model at tissue collection, perhaps reflecting end-stage disease (**Supplementary Fig. 6c**). Nonetheless, levels of fibrosis and inflammation genes were robustly inhibited by either X203 or X209 therapy in this third model (**Supplementary Fig. 6d,e**).

### Neutralisation of IL-11 signalling reverses hepatic fibrosis

While inhibition of IL-11 signalling in mice on HFMCD diet did not reverse total hepatic collagen protein content, there was reversal of *Col1a1*, *Timp1*, and *Tgfβ1* RNA expression (**Supplementary Fig. 5g**). To test more fully, if IL-11 inhibition can reverse the fibrotic phenotype beyond the RNA level, we first established severe liver fibrosis and then removed the fibrogenic stimulus and started antibody treatment (**Fig. 4a**). Hepatic collagen content was significantly reversed after three weeks of X203 or X209 treatment and even greater reversal was seen at 6 weeks (**Fig. 4b, Supplementary Fig. 7a**). In the absence of the dietary trigger, ERK activation spontaneously regressed, which was accelerated by X203 or X209-treatment. Notably, collagen content remained unchanged in IgG control-treated animals for the duration of the experiment (**Fig. 4c, Supplementary Fig. 7b**). As such, anti-IL-11 therapies reverse liver fibrosis but this effect is limited in the context of active and severe steatosis where combination therapy with an anti-steatotic may be more effective.

**Figure 4.**
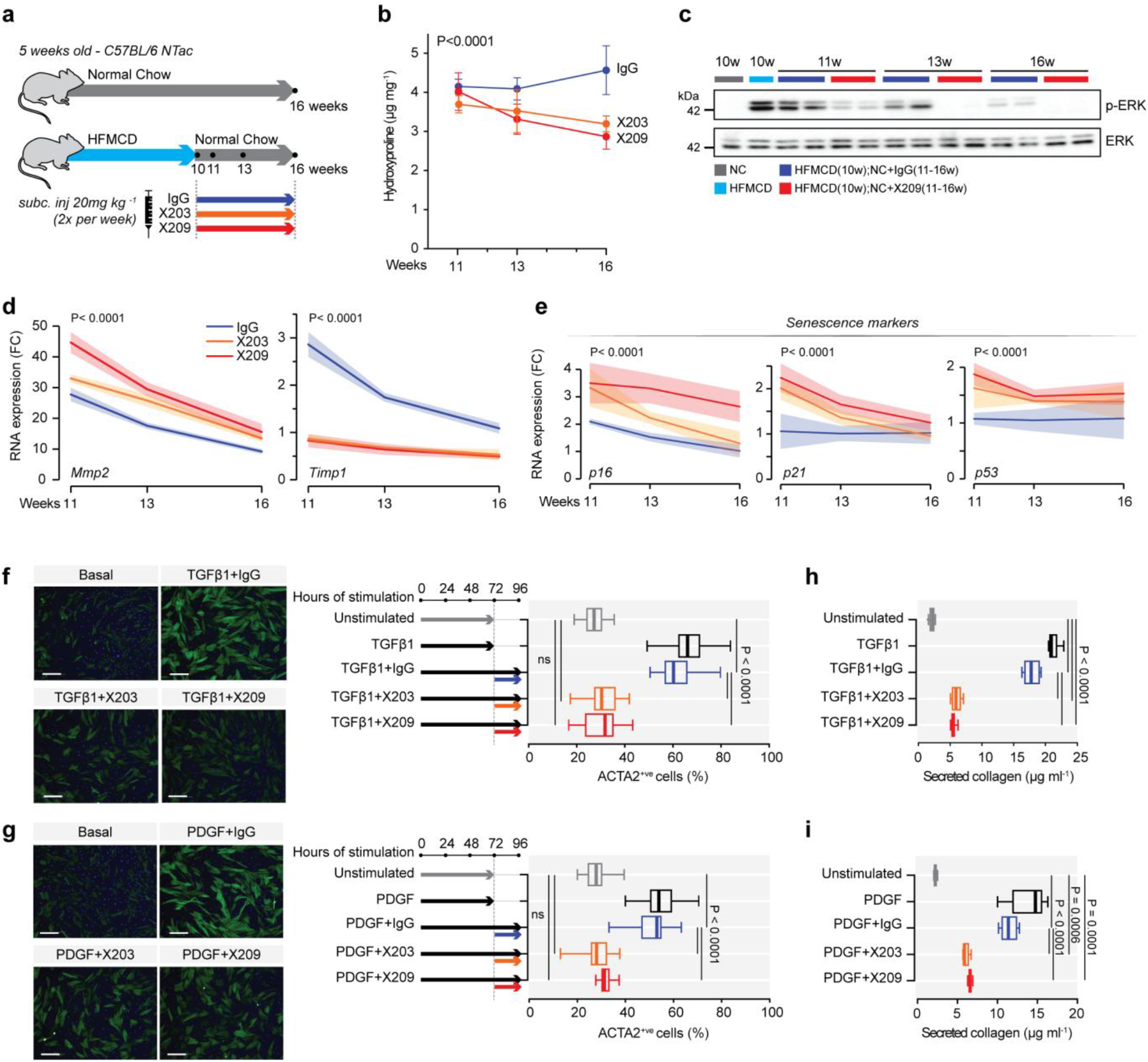
Therapeutic inhibition of Il-11 signalling reverses HSC transformation and liver fibrosis. **a**, Schematic showing reversal experiment with X203 or X209. Fibrosis was established by feeding mice the NASH diet for 10 weeks and then replacing this with normal chow (NC) and initiating antibody (X203 and X209) therapy. Mice were euthanised at the indicated time points. **b-e**, Data for therapy experiments as shown in **4a**. **b**, Total liver hydroxyproline content, **c**, western blots of hepatic ERK activation status, **d**, relative mRNA expression of *Mmp2* and *Timp1*, and **e**, senescence markers at 1-, 3-, 6-weeks after X203, X209, or IgG treatment (n≥3/group). **f, g**, Automated fluorescence quantification and representative fluorescence images of ACTA2^+ve^ immunostaining (scale bars, 200 µm) following incubation without stimulus (-), (**f**) with TGFβ1, (**g**) with PDGF, either prior to or after the addition of X203, X209, or IgG. **h, i**, The amount of collagen secreted by HSCs stimulated with (**h**) TGFβ1 or (**i**) PDGF either prior to or after the addition of IgG, X203, or X209 (n=5/group). **f-i**, TGFβ1 (5 ng ml^-1^), PDGF (20 ng ml^-1)^, IgG, X203 and, X209 (2 µg ml^-1^). **b**, Data are shown as mean ± s.d; **d, e**, data are represented as line chart (mean) and transparencies indicate s.d; **f-i**, data are shown as box-and-whisker with median (middle line), 25th–75th percentiles (box) and min-max percentiles (whiskers). **b, d, e**, Two-way ANOVA; **f-i**, Two-tailed, Tukey-corrected Student’s *t*-test. FC: fold change; NC: normal chow; HFMCD: high fat methionine- and choline-deficient.

Regression of liver fibrosis is associated with lower TIMP and higher MMP levels, which promotes favorable matrix remodelling^3,21^. In keeping with this, X203 or X209 treated mice rapidly exhibited strong upregulation of *Mmp2* and marked downregulation of *Timp1* (**Fig. 4d**). Reversal of hepatic fibrosis is favoured when transformed HSCs undergo apoptosis^22^, senescence^23,24^ or reversion to an inactive, ACTA2^-ve^ cellular state^25^. We examined these potential mechanisms and found decreased *Acta2*, increased senescence markers (*p21*, *p16*, and *p53*) but no change in apoptosis factors with anti-IL-11 therapies (**Fig. 4e, Supplementary Fig. 7c,d**). To check directly if IL-11 signalling is required to maintain HSCs in a transformed state, we stimulated HSCs with TGFβ1 or PDGF and then inhibited IL-11 signalling in the presence of ongoing stimulation. Within 24 h of IL-11 inhibition, the percentage of ACTA2^+ve^ cells and the amount of secreted collagen were reversed to near baseline levels, as was ERK activity (**Fig. 4f-i, Supplementary Fig. 7e,f**). These data show that inhibition of IL-11-dependent HSC transformation causes HSC senescence/reversion and favorable matrix remodelling leading to fibrosis regression.

### Blocking IL-11 signalling inhibits liver inflammation in NASH

Beyond their role in liver fibrosis, HSCs have a central role in hepatic inflammation through the secretion and paracrine activity of pro-inflammatory cytokines and chemokines^3,8,26,27^. We profiled inflammatory gene expression in NASH livers from *Il11ra1^-/-^*, X203- or X209-treated mice and observed consistently lower levels of *TNF*±, *CCL2* and *CCL5* with IL-11 loss-of-function when compared to experimental controls (**Supplementary Fig. 8a,b**). We determined if these effects on inflammation *in vivo* were related to the action of IL-11 on HSCs and found that IL-11 stimulated HSC secretion of CCL2 whereas IL-11 inhibition blocked CCL2 secretion (**Supplementary Fig. 8c)**. This reveals an unappreciated role for IL-11 in stromal immunity and shows that IL-11 neutralisation inhibits paracrine effects of pro-inflammatory factors secreted from HSCs on other cells in the hepatic niche^3,8,26,27^.

In the HFMCD diet model of NASH, inflammation peaks at six weeks and is then followed by a phase of severe fibrosis (**Supplementary Fig. 8d**). We inhibited IL-11 signalling early during steatohepatitis and found that livers of X203- and X209-treated mice were strikingly less steatotic and had lesser ERK activation (**Fig. 5a,b, Supplementary Fig. 8e-g**). At the molecular level, there was a significant reduction in triglyceride content and lipid droplets in hepatocytes of X203- and X209-treated mice were not apparent (**Fig. 5c,d, Supplementary Fig. 8h)**. HFMCD diet induces marked steatohepatitis and liver damage after one week (ALT>700 U L^-1^), which was reversed in a dose-dependent manner to near normal after three weeks of either X203 or X209 treatment (**Fig. 5e,f, Supplementary Fig. 8i**). As expected, X203 or X209 treated mice did not develop fibrosis during the experiment, reaffirming the strong anti-fibrotic effects associated with inhibition of IL-11 signaling (**Fig. 5c,g, Supplementary Fig. 8h,j,k)**.

**Figure 5:**
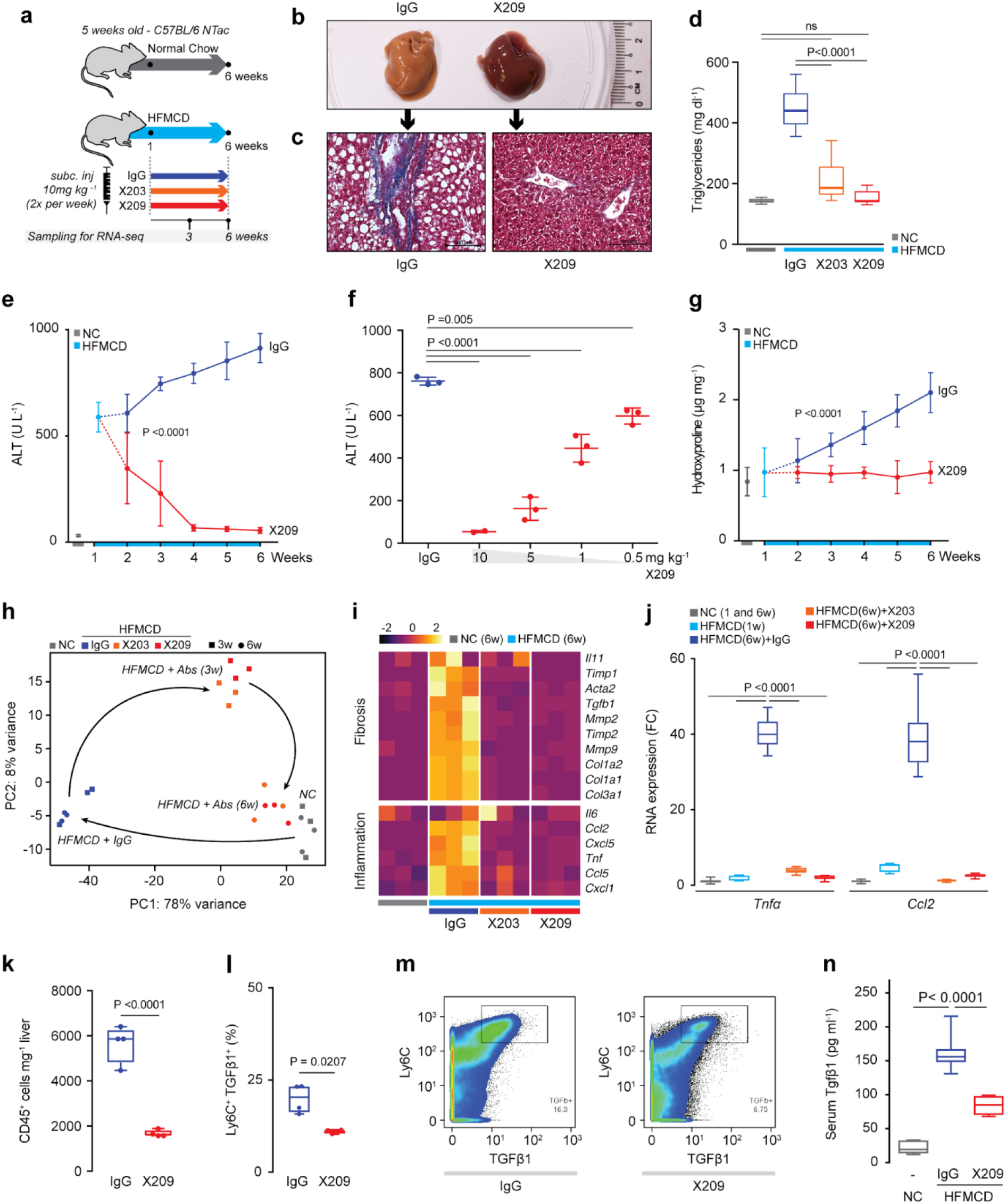
Neutralisation of Il-11 signalling reverses liver damage in early stage NASH. **a**, Schematic of the anti-IL-11 therapy experiment early on in the HFMCD diet NASH model. Antibody treatments were started 1 week after the start of NASH diet when X209, X203, or IgG control (10 mg kg^-1^, twice a week) were administered intraperitoneally for 5 weeks. **b-n**, Data for experiments as shown in **5a**. **b**, Representative gross liver images and **c**, representative Masson’s Trichrome stained images of livers after 5 weeks of IgG or X209 treatments. **d**, Hepatic triglyceride levels (NC, n=5; IgG, n=14; X203, n=10; X209, n=8). **e**, Serum ALT levels (n≥5/group, the values of NC and HFMCD 1 week are the same as those used in **2c**). **f**, Dose-dependent effect of 3-week X209 therapy on reversal of serum ALT levels (n=3/group). **g**, Liver hydroxyproline content of X209- or control IgG-treated mice (n≥5/group, the values of NC and HFMCD 1 week are the same as those used in **Fig. 2b**). **h**, Principal component analysis (PCA) plot of liver gene expression in mice on NC or HFMCD in the presence of IgG, X203 or X209 antibodies for the times shown in **5a** (RNA-seq, n=3/group). Arrows depict the transitions from normal gene expression (NC) to most perturbed gene expression in NASH (HFMCD+IgG), to intermediately restored gene expression (HFMCD+Abs (3w)), to normalised gene expression (HFMCD+Abs(6w)) **i**, Differential expression heatmap of pro-fibrotic and pro-inflammatory genes Z-scores (Transcripts Per Million mapped reads, TPM). **j**, *Tnf*α and *Ccl2* mRNA expression by qPCR (NC, n=9; HFMCD 1 week, n=7; IgG, n=14; X203, n=10; X209, n=8). **k**, Liver CD45^+ve^ immune cell numbers, **l**, Ly6C^+ve^ TGFβ1^+ve^ cells in the total CD45^+ve^ populations and **m**, representative pseudocolor plots illustrating the gating strategy used to detect Ly6C^+ve^ TGFβ1^+ve^ cells. **k-m**, (n=4/group). **n** Serum TGFβ levels (NC, n=5; IgG, n=14; X203, n=10; X209, n=8). **d, j, k, l, n**, Data are shown as box-and-whisker with median (middle line), 25th–75th percentiles (box) and min-max percentiles (whiskers); **e-g**, data are shown as mean ± s.d. **d, j, n**, Two-tailed, Tukey-corrected Student’s *t*-test; **e, g**, two-way ANOVA; **f**, two-tailed Dunnett’s test; **k, l**, two-tailed Student’s *t*-test. FC: fold change; NC: normal chow; HFMCD: high fat methionine- and choline-deficient.

We next performed RNA-seq to profile globally the effects of IL-11 inhibition during steatohepatitis. Unsupervised analyses of these data showed that antibody treatment almost completely reverses the pathological RNA expression signature induced by HFMCD diet (**Fig. 5h, Supplementary Fig. 9a,b**). Upregulation of pro-fibrotic and pro-inflammatory genes was abolished and lipid metabolism gene expression re-established by anti-IL11RA therapy (**Fig. 5i,j, Supplementary Fig. 9c-e**). Unbiased Gene Set Enrichment Analyses confirmed the reversion of HFMCD diet-induced changes in metabolic and inflammatory transcriptional signatures (**Supplementary Fig. 10**).

Resident macrophages and infiltrating monocytes are important for NASH pathogenesis and a major source of TGFβ1 during disease progression^28^. We examined inflammatory cell populations in the liver during steatohepatitis and observed fewer immune cells in general in X209-treated livers and a specific reduction in Ly6C^+ve^TGFβ1^+ve^ cells (**Fig. 5k-m**). TGFβ1 is a major determinant of fibrosis in NASH and can be produced by Kupffer, HSCs and other cells in the liver^5^. Circulating TGFβ1 levels were elevated by HFMCD diet but reduced by X209 therapy, which shows that anti-IL11RA therapy is disease-modifying (**Fig. 5n**).

## Discussion

Recognition of HSCs as the major source for myofibroblasts in the liver^2^ prioritizes their transformation as a specific and fundamental target in fibrotic liver diseases. We have previously identified an important function of IL-11 for cardiac and renal fibroblast-to-myofibroblast transformation^10^. We reveal here that IL-11 has non-redundant signalling activity required for HSC activation and transformation, which is positioned at a decisive intersection of several pathogenic pathways. Our findings show that non-canonical IL-11 signalling is an overlooked and cardinal process for myofibroblast generation from both fibroblasts and HSCs, and likely pericytes and other cell types, and confirm a key role for ERK signaling in hepatic fibrosis^29^.

The multi-faceted pathobiology of HSCs touches upon many aspects of liver disease: fibrosis, metabolism, immunoregulation and secretion of paracrine factors in the hepatic niche^3^. Confirming the central role of IL-11 for HSC pathobiology, our first-in-class IL-11 neutralising treatments show disease-modifying therapeutic impact beyond anti-fibrotic effects alone. Inhibition of IL-11 signaling prevents inflammation and steatosis and can reverse liver fibrosis and hepatocyte damage. Unlike steatosis, fibrosis in NASH predicts clinical endpoints and anti-fibrotic IL-11 blocking therapies may offer benefits over drugs that primarily target liver metabolism^1^.

While earlier publications suggest IL-11 is anti-inflammatory in liver^12,13^, these studies use high-dose recombinant human IL-11 in the mouse, where effects can be non-specific^10^. In contrast, we show here that the biological effect of endogenous Il-11 at physiological levels is the opposite: HSC-immune cell crosstalk and activation is IL-11 dependent and inhibition of IL-11 is anti-inflammatory and cytoprotective. Our study demonstrates robust modulation of the immune response by targeting stromal cells through IL-11 inhibition, which was unexpected. This may have implications for other fibro-inflammatory processes where stromal and immune cell functions are closely interlinked, as in tumour microenvironments^30,31^ and autoimmune diseases^32,33^.

Human^34^ and mouse^17^ knockouts of *IL11RA* have mild developmental abnormalities of the skull but are otherwise healthy and IL-11 appears largely redundant in adult mammals. This provides compelling genetic safety data for IL-11 as a viable drug target and we suggest that IL-11 neutralising therapies should be evaluated in NASH. This is the first study to demonstrate a role for IL-11 in HSC biology, NASH or stromal immunity and lays the groundwork for future studies to dissect fully the effects of IL-11 signaling in the liver. We believe this presents an exciting opportunity and that our findings may have broad implications across tissues and diseases.

## Acknowledgements

The authors would like to acknowledge the technical expertise and support of N.S-J.Ko, B.L.George, M. Wang, and NGS Team at NHCS. The research was supported by the National Medical Research Council (NMRC) Singapore STaR awards to S.A.C. (NMRC/STaR/0029/2017), the NMRC Central Grant to the NHCS, Goh Foundation, Tanoto Foundation and a grant from the Fondation Leducq. A.A.W. is supported by the NMRC YIRG (NMRC/OFYIRG/0053/2017). B.K.S. is supported by the NMRC YIRG (NMRC/OFYIRG/0002/2016). P.M.Y. is supported by NMRC/CSA/0054/2013 and NMRC/CIRG/1457/2016.

## Author contributions

A.A.W., B.K.S., S.S., and S.A.C. conceived and designed the study. A.A.W., B.K.S., S.V., J.R.D., B.N., J.T., and M.T. performed *in vitro* cell culture, cell biology and molecular biology experiments. A.A.W., B.K.S., J.T., M.T., A.R., M.S., E.B., and R.A.S. performed *in vivo* gain- and loss-of function mouse studies. N.G-C. and S.M.E. performed gain-of function studies on *Col1a1-GFP* mice. A.A.W., W.W.L., and S.Y.L, performed histology analysis. A.A.W and S.V. performed *in vitro* antibody screening. G.D., S.P.C., and S.S performed computational analysis. B.S.P and S.A. performed CyTOF. A.A.W., B.K.S, E.A., G.D., B.N., R.A.S., P.M.Y., S.S., and S.A.C analyzed the data. A.A.W., E.A., S.S., and S.A.C prepared the manuscript with input from co-authors.

## Supplementary Figures

**Supplementary Figure 1.**
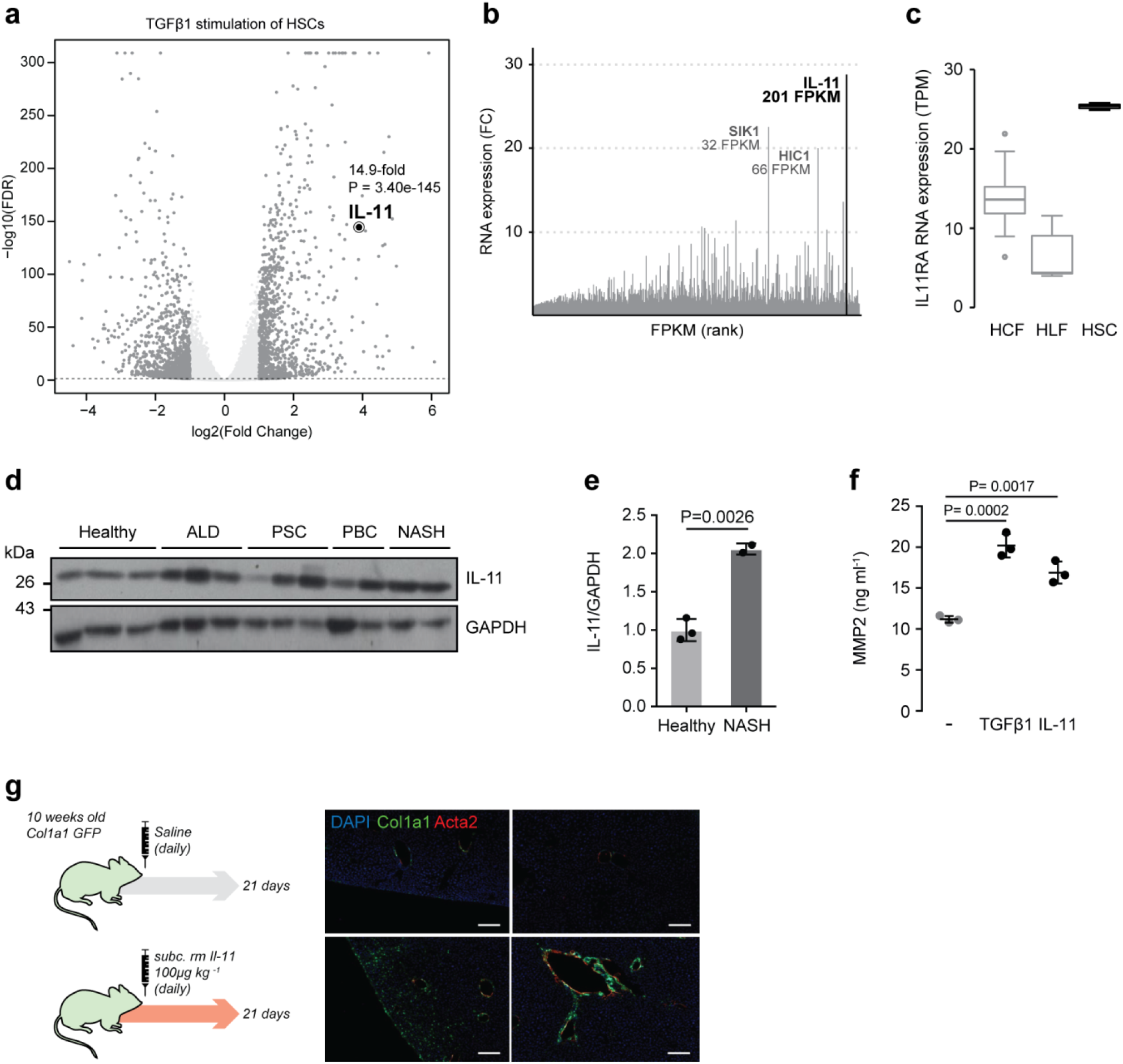
HSCs secrete and respond to IL-11. **a**, Genome-wide changes in RNA expression in HSCs (n=3) after TGFβ1 stimulation (5 ng ml^-1^, 24 h). **b**, Stiffness-induced RNA upregulation in humans HSCs (RNA-seq^14^), genes are ranked according to fragments per kilobase million (FPKM), *IL-11* upregulation is the most highly upregulated gene genome wide. **c**, IL11RA transcripts in human cardiac fibroblasts (HCF), human lung fibroblasts (HLF), and human HSC (TPM, Transcript per millions). Data are shown as box-and-whisker with median (middle line), 25th–75th percentiles (box) and min-max percentiles (whiskers). **d**, Western blots and **e**, densitometry of IL-11 and GAPDH in human liver samples of healthy individuals and patients suffering from alcoholic liver disease (ALD), primary sclerosing cholangitis (PSC), primary biliary cirrhosis (PBC), and non-alcoholic steatohepatitis (NASH). Data are shown as scatter plot with bar, mean ± s.d; two-tailed Student’s *t*-test. **f**, MMP-2 concentration in the supernatant of HSC (n=3/group) without stimulus (-), with TGFβ1 or IL-11 (5 ng ml^-1^, 24 h) by ELISA. Data are represented as mean ± s.d; two-tailed Dunnett’s test. **g**, Schematic and representative fluorescence images GFP^+ve^ cells of *Col1a1-GFP* mice injected daily with either rmIl-11 (100 μg kg^-1^) or saline. Sections were immunostained for Acta2 and counterstained with DAPI (scale bars, 200 μm). FC: fold change.

**Supplementary Figure 2.**
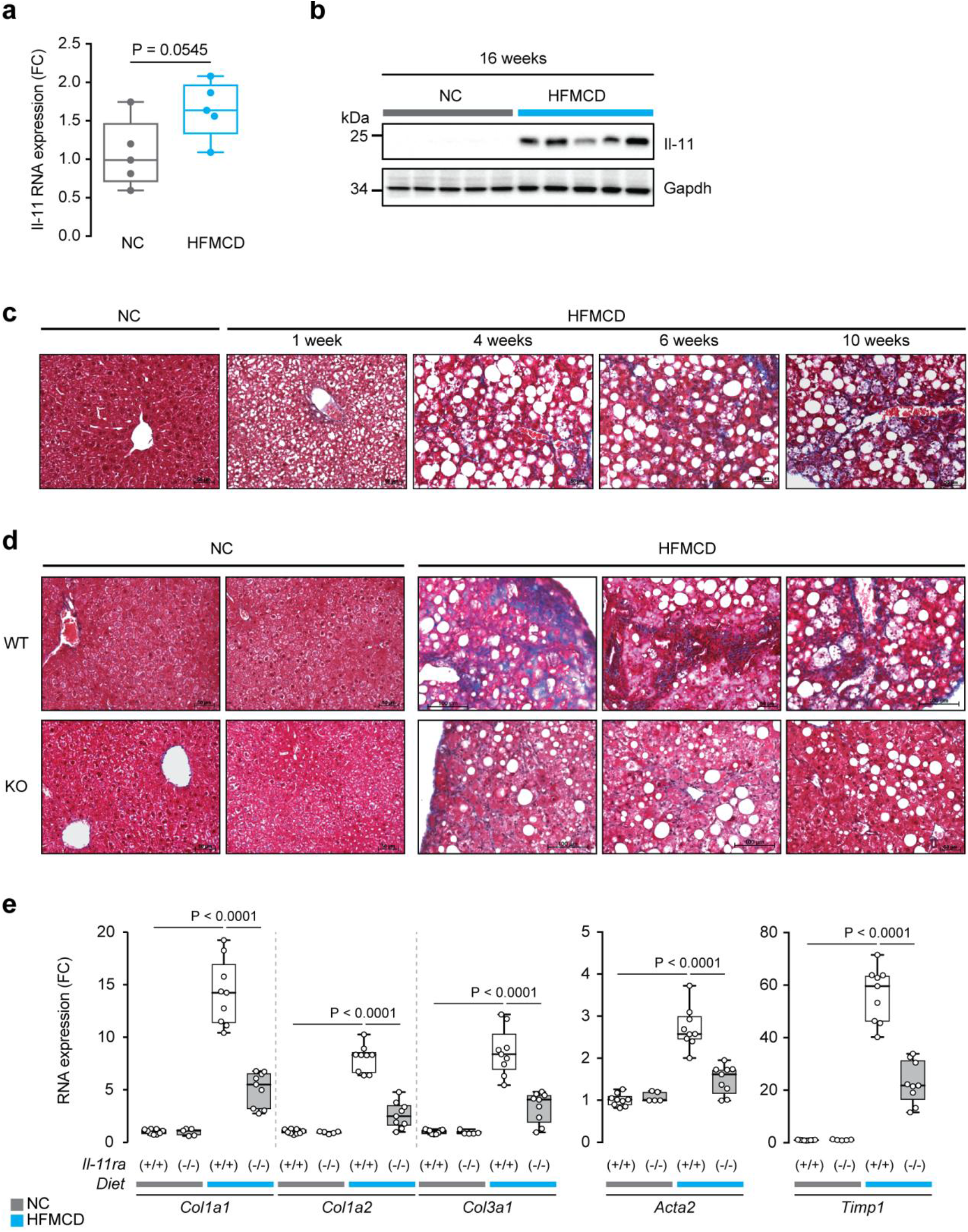
Genetic inhibition of Il-11 signalling reduces hepatic fibrosis. **a, b**, Effects of 16 weeks of HFMCD diet as compared to NC diet on hepatic (**a**) *Il-11* mRNA and (**b**) Il-11 protein levels. For (**a**) and (**b**) RNA and protein were extracted from the same mice (n=5/group). **c**, Representative Masson’s Trichrome images of liver sections from mice fed with NC or HFMCD diet for the indicated treatment duration. **d**, Representative Masson’s Trichrome images and **e**, relative RNA expression level of *Acta2*, *Col1a1*, *Col1a2*, and *Col3a1* in livers of *Il11ra^+/+^* (WT) and *Il11ra^-/-^* (KO) mice after 10 weeks of HFMCD diet. **e**, NC WT, n=9; HFMCD WT, n=9, NC KO, n=5; HFMCD KO, n=9. **a, e**, Data are shown as box-and-whisker with median (middle line), 25th–75th percentiles (box) and min-max percentiles (whiskers). **a**, Two-tailed Student’s *t*-test; **e**, Sidak-corrected Student’s *t*-test. FC: fold change; NC: normal chow; HFMCD: high fat methionine- and choline-deficient.

**Supplementary Figure 3.**
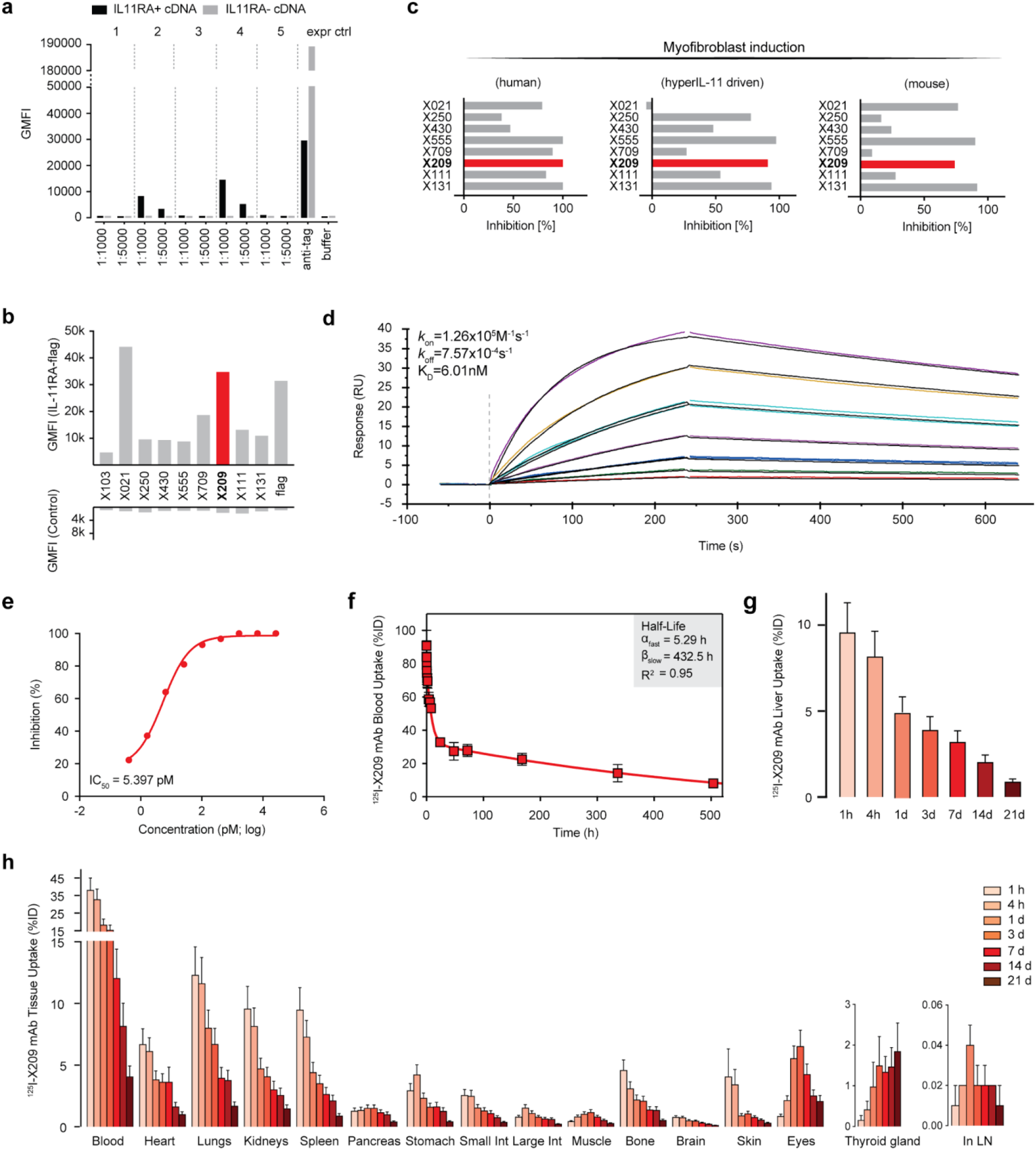
Development of a neutralizing anti-IL-11RA monoclonal antibody. **a**, Sera of 5 mice after genetic immunization with human IL11RA. Sera of five animals tested with HEK cells transiently transfected with an *IL11RA-flag* or control cDNA vector, incubated with a goat anti-mouse fluorescent antibody (10 µg ml^-1^). Cells were then analysed by flow cytometry. Signal is geometric mean of the relative fluorescence (GMFI) as measured by flow cytometry. **b**, Supernatants of early stage hybridoma cultures on transfected cells. **c**, Inhibition of ACTA2^+ve^ cell transformation of TGFβ1-(left), hyperIL-11-(middle) stimulated human atrial fibroblasts and TGFβ1-(right) stimulated mouse atrial fibroblasts with purified mouse monoclonal anti-IL11RA candidates (6 µg ml^-1^). **d**, X209 interactions with IL11RA as determined by SPR (1:1 Langmuir). **e**, Dose-response curve and IC_50_ value of X209 (61 pg ml^-1^ to 4 µg ml^-1^; 4-fold dilution) in inhibiting MMP2 secretion by HSCs stimulated with TGFβ1. **c, e**, TGFβ1 (5 ng ml^-1^), Hyper IL-11 (0.2 ng ml^-1^); 24 h. **f**, Blood pharmacokinetics of ^125^I-X209 in mice (n=5). Result was fitted (R^2^=0.92) to a two-phase exponential decay model. **g, h**, Percentage of ^125^I-X209 uptake by (**g**) liver (n=5) and (**h**) other organs at the indicated time points, following retro-orbital injection. **f-h**, Data are represented as mean + s.d.

**Supplementary Figure 4.**
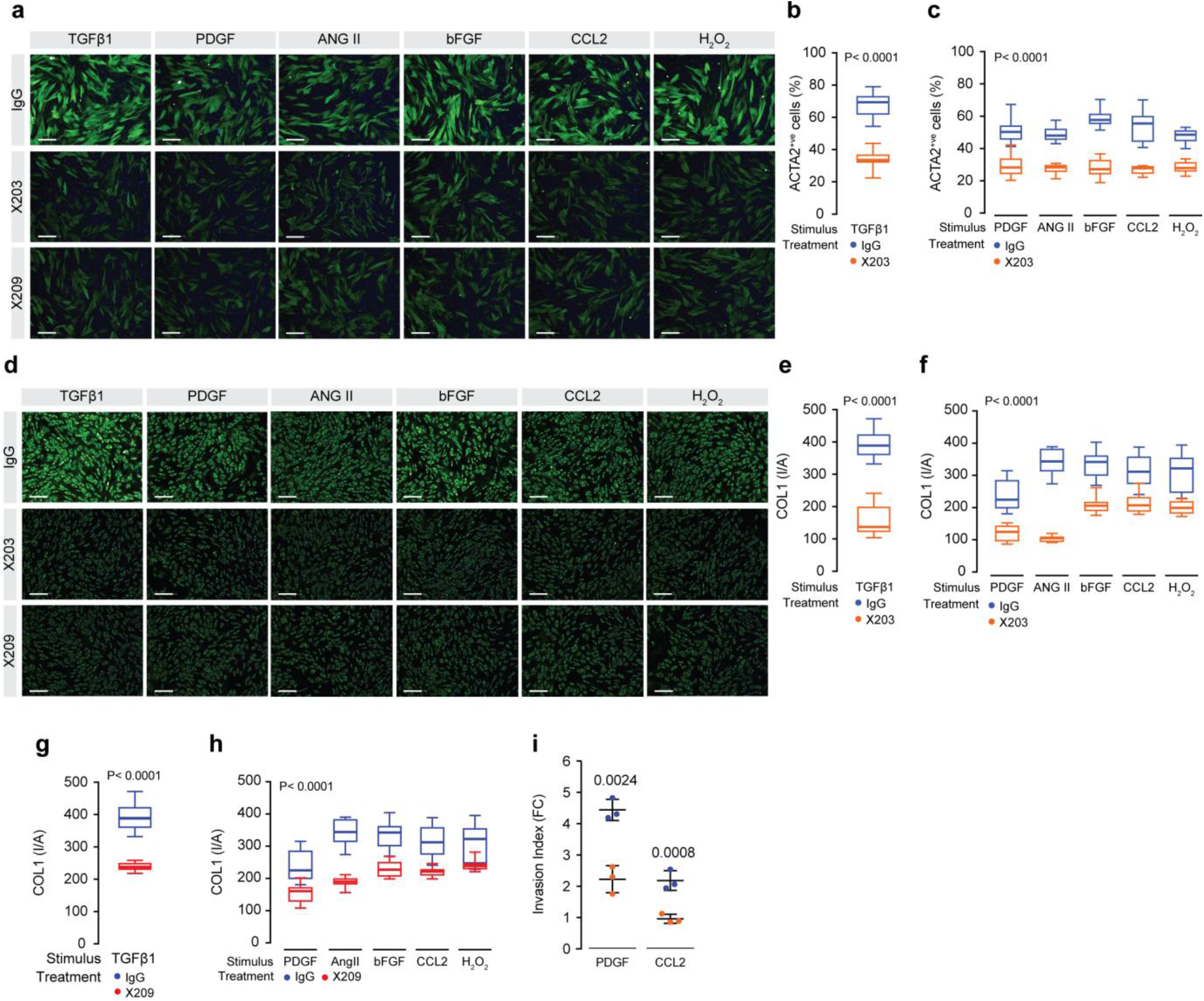
Neutralizing anti-IL-11 and anti-IL11RA antibodies prevent HSC-to-myofibroblasts transformation. **a-h**, (**a, d**) Representative fluorescence images (scale bars, 200 µm) and (**b,c,e-h**) quantification of (**a-c**) ACTA2^+ve^ cells and (**d-h**) Collagen 1 immunostaining of HSCs treated with (**a,b,d,e,g**) TGFβ1 and other (**a,c,d,f,h**) NASH factors in the presence of IgG control, X203, or X209 for 24 h. **i**, Effects of X203 on TGFβ1- and CCL2-induced matrigel invasion of HSCs for 48h (n=3). **a-i**, TGFβ1 (5 ng ml^-1^), Hyper IL-11 (0.2 ng ml^-1^), PDGF (20 ng ml^-1^), AngII (100 nM), bFGF (10 ng ml^−1^), CCL2 (5 ng ml^-1^), H_2_ O_2_ (0.2 mM), IgG, X203 and X209 (2 μg ml^-1^). **b, c, e-i**, Two-tailed Dunnett’s test. **b, c, i**, The values of IgG are the same as those shown in **Fig. 2h,j,k** respectively. The values of IgG for **g** and **h** are the same as those shown in **c** and **f**, respectively. **b, c, e-h**, Data are shown as box-and-whisker with median (middle line), 25th–75th percentiles (box) and min-max percentiles (whiskers); **i**, data are represented as mean ± s.d. FC: fold change; I/A: intensity/area.

**Supplementary Figure 5.**
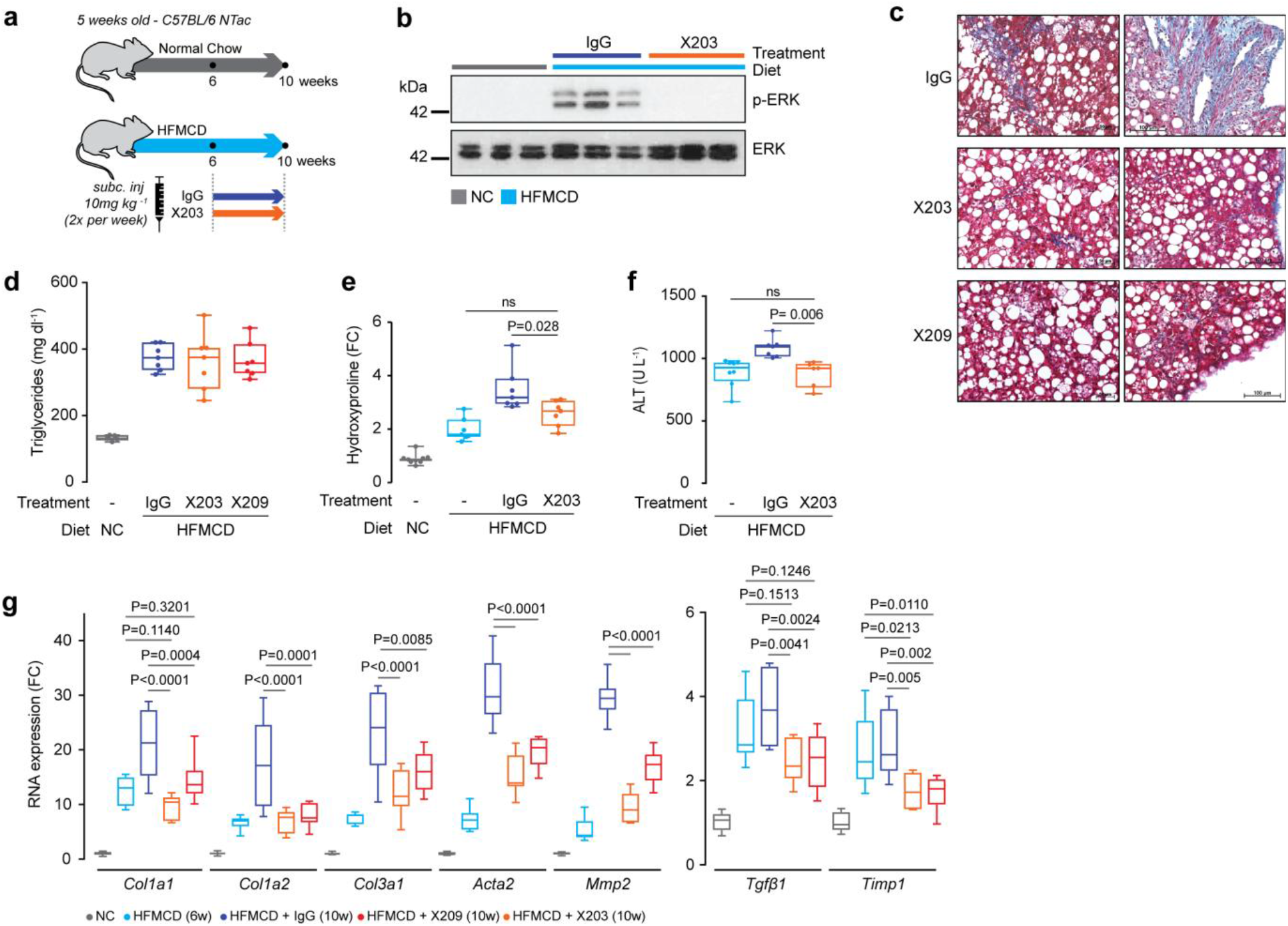
Neutralizing anti-IL-11 and anti-IL11RA antibodies inhibit hepatic fibrosis and liver damage. **a**, Schematic of therapeutic dosing regimen of X203 in HFMCD fed mice. X203 or IgG (10 mg kg^-1^, twice a week) were administered for 4 weeks, starting from week 6 of the NASH diet. Livers and serum were collected at week 10. **b-g**, Data for therapeutic dosing experiments as shown in **Fig. 3a** and **Supplementary Data Fig. 5a**. **b**, Western blots of liver ERK activation, **c**, representative histological images (Masson’s Trichrome staining) of liver sections, **d**, liver triglyceride content, **e**, relative liver hydroxyproline collagen content, and **f**, serum ALT levels from IgG- and X203-treated mice. **d**, NC, n=5; IgG, n=7; X203, n=7, X209, n=7. **e**, The values of NC and HFMCD 6 weeks are the same as those used in **Fig. 2b**; the values of IgG are the same as those used in **Fig. 3c**; X203, n=7. **f**, The values of HFMCD 6 weeks are the same as those used in **Fig. 2c**; the values of IgG are the same as those used in **Fig. 3e**; X203, n=6. **g**, Expression levels of liver pro-fibrotic genes (NC, n=9; HFMCD 6 weeks, n=8; IgG, n=8; X203, n=7; X209, n=9). **d-g**, Data are shown as box-and-whisker with median (middle line), 25th–75th percentiles (box) and min-max percentiles (whiskers); two-tailed, Tukey-corrected Student’s *t*-test. FC: fold change; NC: normal chow; HFMCD: high fat methionine- and choline-deficient.

**Supplementary Figure 6.**
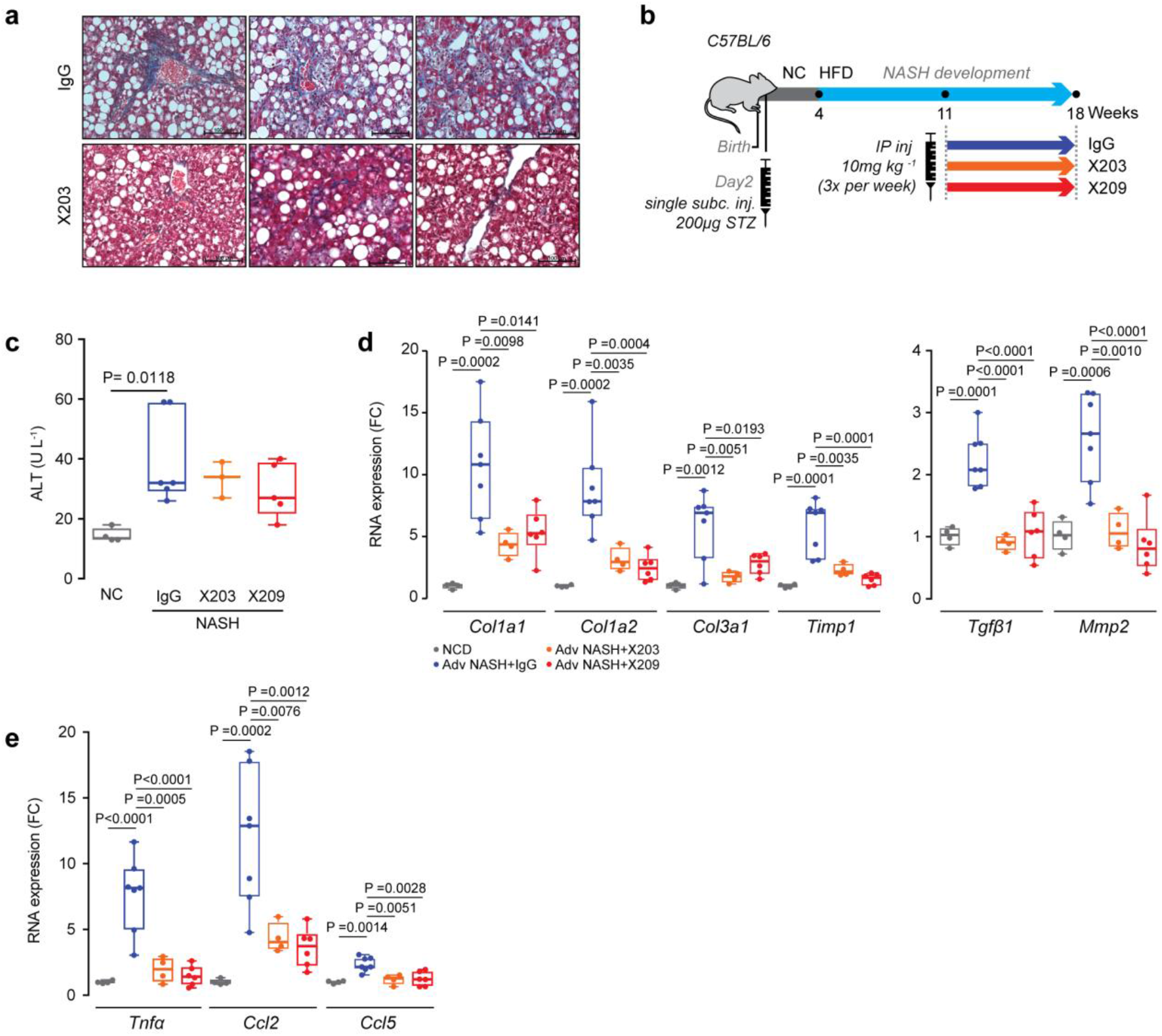
Neutralizing anti-IL-11 and anti-IL11RA antibodies reduce hepatic fibrosis and hepatic inflammation in additional NASH models. **a**, Representative Masson’s Trichrome images of livers from *db/db* mice treated with X203 or IgG. **b**, Schematic representation of the STAM^™^ model. Mice were injected with 200 µg of Streptozotocin (STZ) 2 days after birth followed by feeding with high fat diet (HFD) at 4-week of age to develop NASH. IgG, X203, and X209 were intraperitoneally injected 3x/week at a dosage of 10 mg kg^-1^ for 7 weeks, starting at 11-weeks of age. **c**, Serum ALT levels of STAM^™^ mice (NC, n=4; IgG, n=6; X203, n=3; X209, n=5). **d, e**, Relative liver mRNA expression levels of fibrosis (**d**) and inflammation (**e**) genes in STAM^*™*^ mice treated with X203 or X209 (NC, n=4; IgG, n=7; X203, n=4; X209, n=6). **c-e**, Data are shown as box-and-whisker with median (middle line), 25th–75th percentiles (box) and min-max percentiles (whiskers); two-tailed, Tukey-corrected Student’s *t*-test. FC: fold change; NC: normal chow; HFD: high fat diet.

**Supplementary Figure 7.**
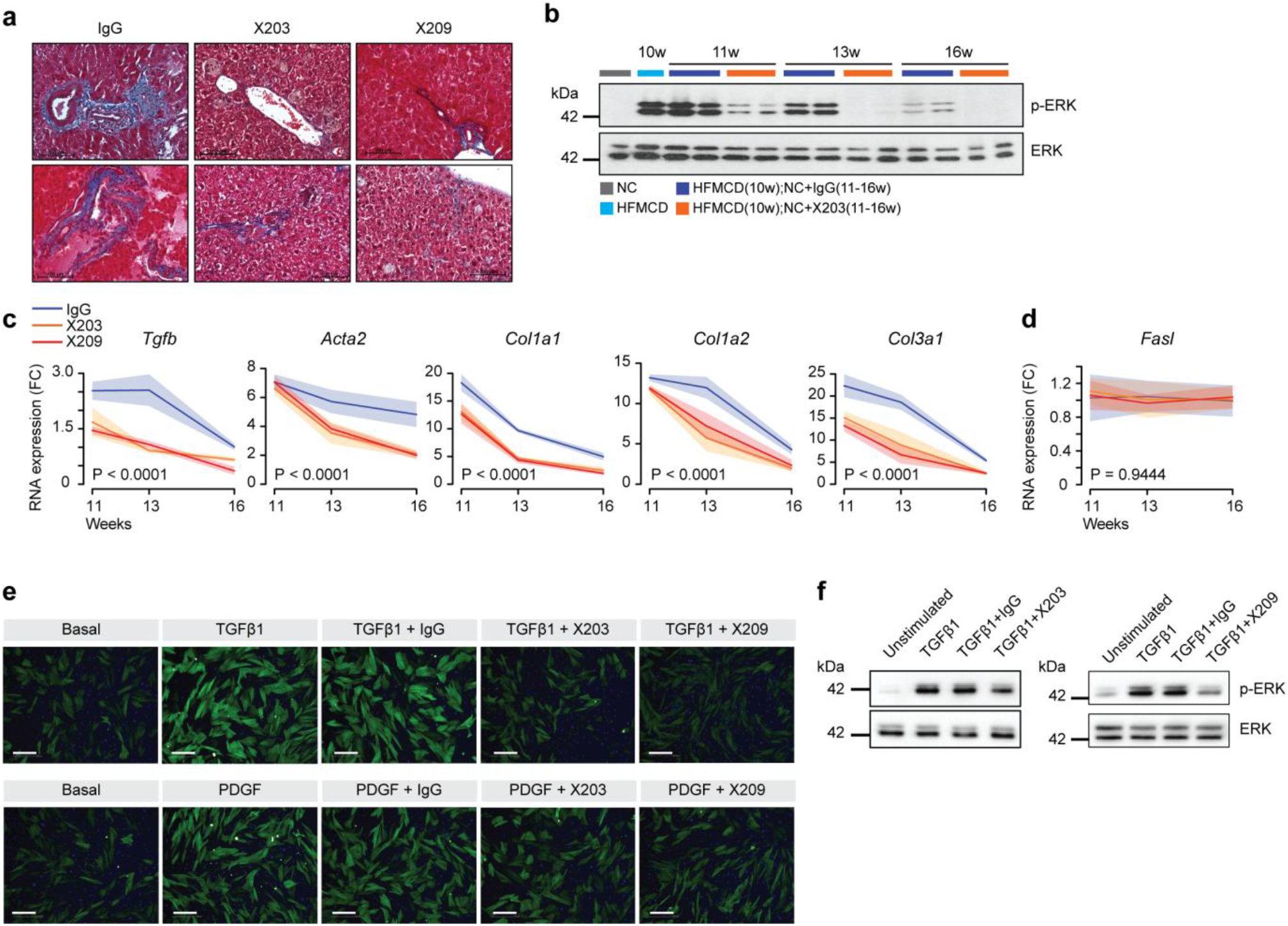
Neutralizing anti-IL-11 and anti-IL11RA antibodies reverse hepatic fibrosis. **a**, Representative Masson’s Trichrome staining of livers from mice treated with IgG, X203, or X209 for 6 weeks as shown in **Fig. 4a**. **b**, Western blots of hepatic ERK activation status. **c, d**, Relative mRNA expression of (**c**) fibrosis markers and (**d**) apoptosis marker (*Fasl*) at 1-, 3-, 6-weeks after X203, X209, or IgG treatments (n≥3/group); two-way ANOVA. **e,f**, Data from reversal of HSC transformation experiments as shown in **Fig. 4f,g**, TGFβ1 (5 ng ml^-1^), PDGF (20 ng ml^-1^), IgG, X203, and X209 (2 μg ml^-1^). **e**, Representative fluorescence images (scale bars, 200 µm) of ACTA2^+ve^ immunostaining following incubation with TGFβ1 or with PDGF either prior to or after addition of X203, X209, or IgG. **f**, Western blots of ERK activation status after X203 and X209 treatment in TGFβ1-treated HSC. **c, d**, Data are represented as line chart (mean) and transparencies indicate s.d. FC: fold change; NC: normal chow; HFMCD: high fat methionine- and choline-deficient.

**Supplementary Figure 8.**
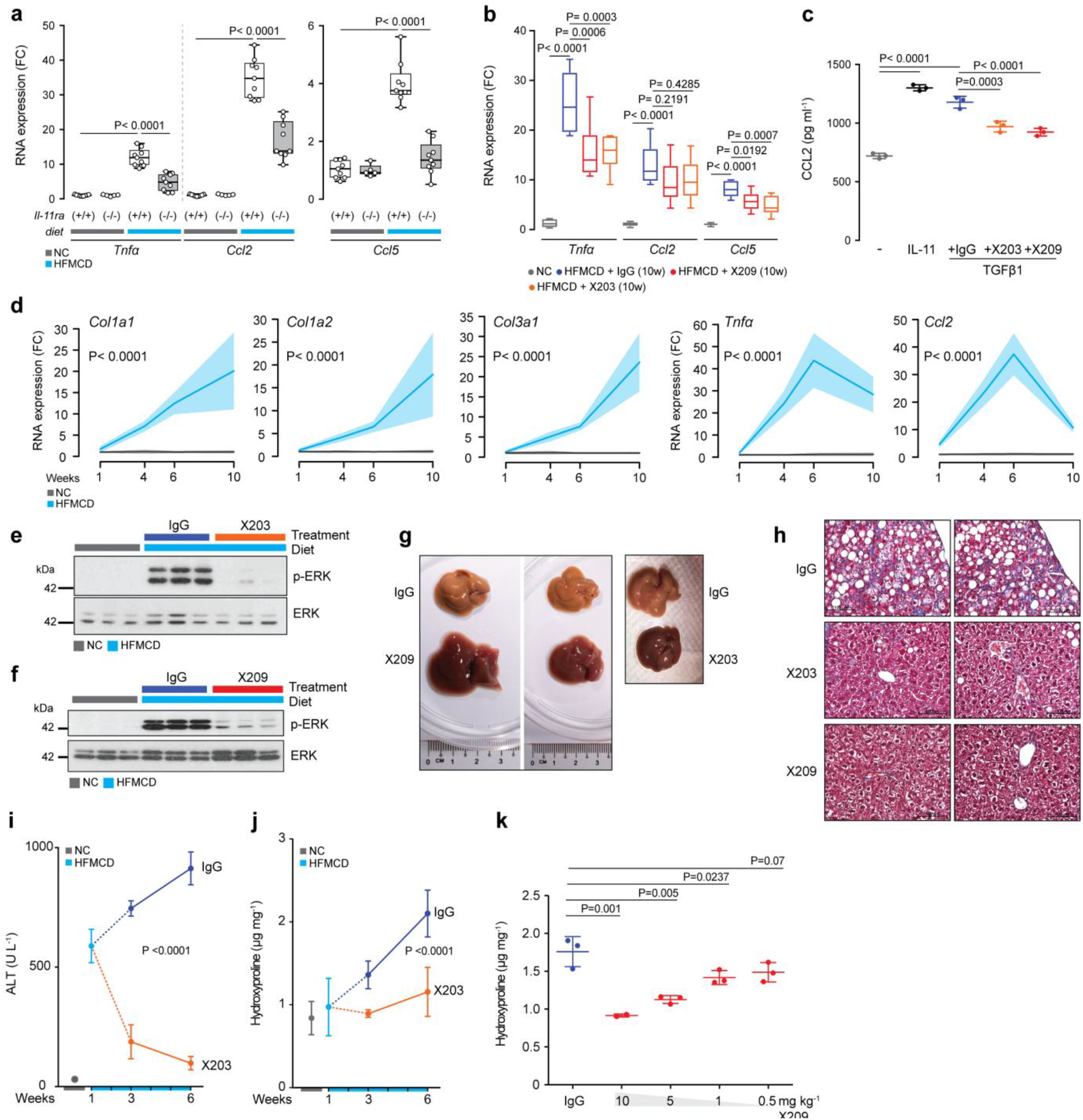
Neutralizing anti-IL-11 and anti-IL11RA antibodies prevent hepatic fibrosis and reduce hepatic inflammation in HFMCD fed mice. **a, b**, Relative mRNA expression of inflammation markers (*Tnf*α, *Ccl2*, and *Ccl5*) from the livers of (**a**) *Il11ra^+/+^* (WT) and *Il11ra^-/-^* (KO) after 10 weeks of HFMCD diet and (**b**) mice injected with X203 or X209 as shown in **Fig. 3a** and **Supplementary Data Fig. 5a**. **a**, NC WT, n=9; HFMCD WT, n=8, NC KO, n=5; HFMCD KO, n=9. **b**, NC, n=9; IgG, n=8; X203, n=7; X209, n=9. **c**, CCL2 in the supernatants of HSCs (n=4/group) without stimulus (-), with IL-11, or with TGFβ1 in the presence of IgG, X203, or X209 by ELISA; IL-11 (5 ng ml^-1^), TGFβ1 (5 ng ml^-1^), IgG, X203, and X209 (2 μg ml^-1^). **d** Relative liver mRNA expression of fibrosis and inflammation markers from mice fed with NC or HFMCD diets. Livers were collected at the indicated time points. The values of NC 1 and 6 week(s) for *Tnf*α and *Ccl2* are the same as those shown in **Fig. 5j**; the values of NC 6 and 10 weeks and HFMCD 6 weeks are the same as those shown in **Supplementary Data Fig. 5g and 8b** (n≥5/group). **e-k**, Data for therapeutic dosing experiments as shown in **Fig. 5a. e-f**, Western blots of hepatic ERK activation status after (**e**) X203 and (**f**) X209 treatments. **g**, Representative gross liver images, **h**, representative Masson’s Trichrome stained images of livers, **i**, serum ALT levels, **j**, liver hydroxyproline content, and **k**, dose dependent effects of X209 on total hydroxyproline content in HFMCD-fed mice (n=3/group). **i**, The values of NC and HFMCD 1 week are the same as those used in **Fig. 2c**, the values of IgG 3 and 6 weeks (2 weeks and 5 weeks treatment, respectively) are the same as those used in **Fig. 5e**. **j**, The values of NC and HFMCD 1 week diets are the same as those used in **Fig. 2b**, the values of IgG 3 and 6 weeks are the same as those used in **Fig. 5g**. **i,j**, X203 3 weeks, n=5; X203 6 weeks, n=10). **a, b**, Data are shown as box-and-whisker with median (middle line), 25th–75th percentiles (box) and min-max percentiles (whiskers); **c, i-k**, data are represented as mean ± s.d; **d**, data are represented as line chart (mean) and transparencies indicate s.d. **a**, Two-tailed, Sidak-corrected Student’s *t*-test; **b, c**, two-tailed, Tukey-corrected Student’s *t*-test; **d, i, j**, two-way ANOVA; **k**, two-tailed Dunnett’s test. FC: fold change; NC: normal chow; HFMCD: high fat methionine- and choline-deficient.

**Supplementary Figure 9.**
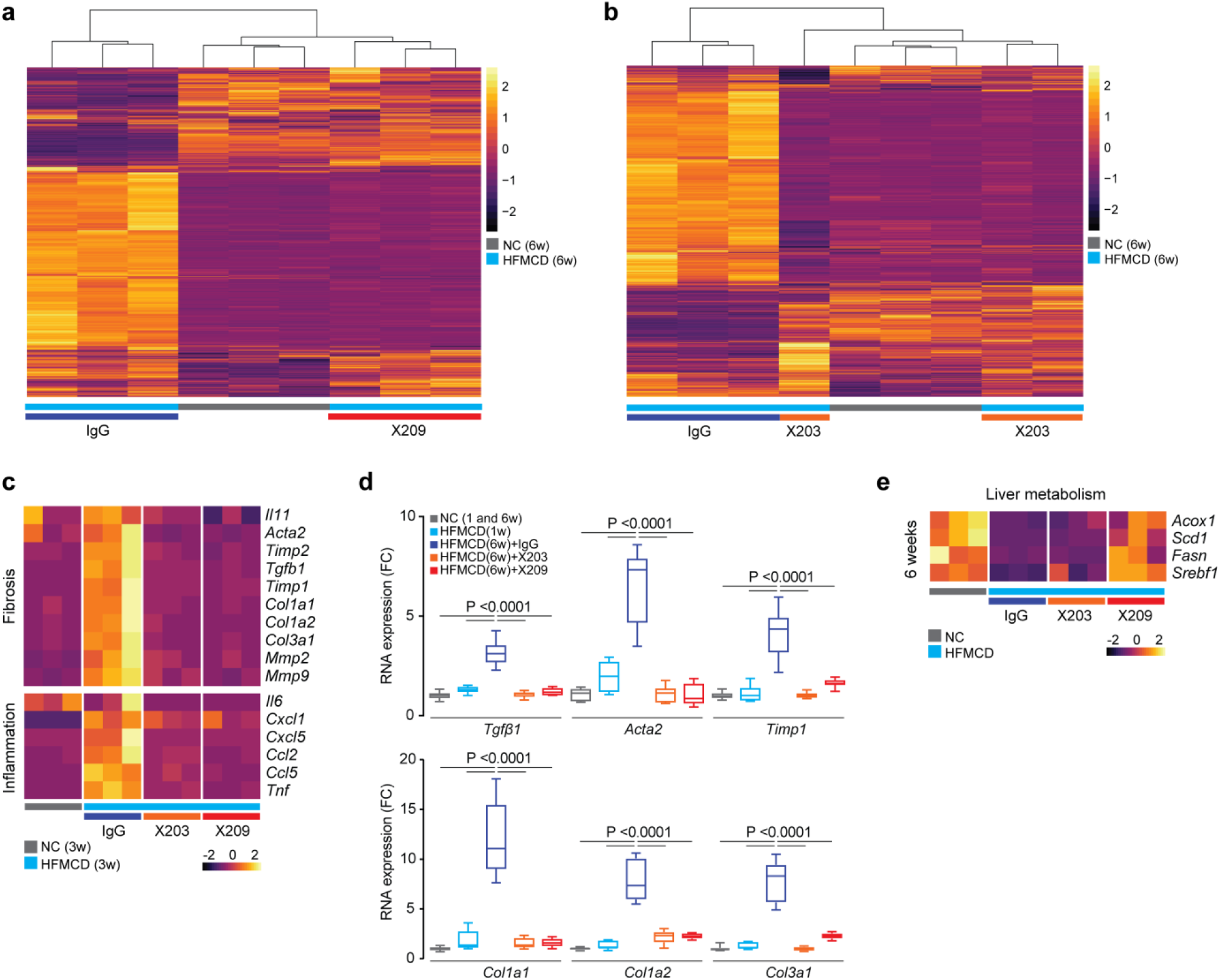
Neutralizing anti-IL-11 or anti-IL11RA antibodies reverse the molecular signature of NASH towards a normal liver profile. **a-e**, Data for RNA-seq and qPCR confirmation for early therapeutic dosing experiments as shown in **Fig 5a. a-b**, Heatmaps showing gene expression levels (Transcripts Per Million mapped reads, TPM) across samples for all genes statistically differentially expressed between IgG and (**a**) X209 or (**b**) X203 treatments. The expression profile for the anti-IL-11 treatments clusters together with the profiles in normal chow (NC), suggesting an almost complete reversal of the transcriptional effect of HFMCD diet. **c**, Heatmaps showing Z-scores of TPM of pro-fibrotic and pro-inflammatory genes indicate that the difference between NC and HFMCD diet were already largely restored by both X203 and X209 within 2 weeks of starting treatment. **d**, Relative RNA expression levels of fibrosis markers after 5 weeks treatment of X203 and X209 by qPCR, which confirms data from RNA-seq. Data are shown as box-and-whisker with median (middle line), 25th–75th percentiles (box) and min-max percentiles (whiskers); two-tailed, Tukey-corrected Student’s *t*-test. The values of NC and HFMCD 1 week for *Col1a1*, *Col1a2*, and *Col3a1* are the same as those shown in **Supplementary Data Fig. 5g, 8d**; NC, n=9; HFMCD 1 week, =7, IgG, n=14; X203, n=10; X209, n=8. **e**, Differential expression heatmap of lipogenesis and β-oxidation genes showing that X209, more so than X203, improved hepatic lipid metabolism as compared to IgG. **a-c, e**, n=3/group. FC: fold change; NC: normal chow; HFMCD: high fat methionine- and choline-deficient.

**Supplementary Figure 10.**
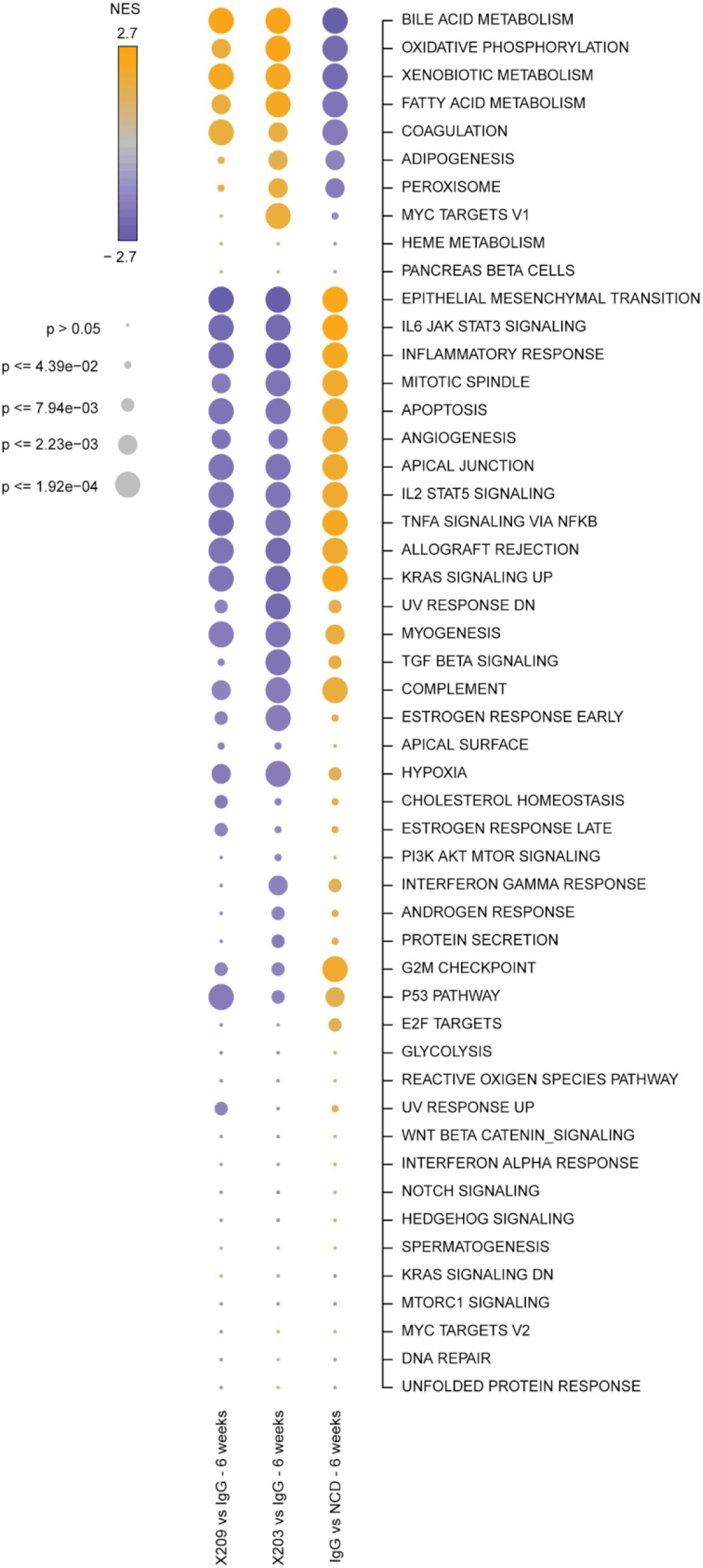
Gene Set Enrichment Analysis of the effects of anti-IL-11 or anti-IL11RA antibody therapy in mice of HFMCD diet as compared to control. Bubblemap showing results of the Gene Set Enrichment Analysis (GSEA) for differentially expressed genes found in every comparison after 6-weeks of NC or HFMCD diet and antibody therapy, as shown. Each dot represents the Normalized Enrichment Score (NES) for the gene set and its FDR-corrected significance level, summarized by colour and size respectively. Gene sets for the enrichment test were selected from the “H - Hallmark” collection in MSigDB.

## Material and Methods

### Animal experiments

All animal procedures were approved and conducted in accordance with the SingHealth Institutional Animal Care and Use Committee (IACUC). All mice were provided food and water *ad libitum*.

#### Mouse models of NASH

##### High fat methionine- and choline-deficient (HFMCD) diet fed mice

Five-week old male C57BL/6N mice were fed methionine- and choline-deficient diet supplemented with 60 kcal% fat (A06071301B, Research Diets); control mice were fed with normal chow (NC, Specialty Feeds). Durations of diet and antibody therapies varied as outlined in the main text.

##### MCD diet fed Leprdb/db mice

Male BKS.Cg-Dock7m+/+LeprdbJ (*db*/*db*) mice were used when they are at 12-weeks of age and at the hepatic steatosis stage. Animals were then fed methionine- and choline-deficient diet (MCD, A02082002BRi, Research Diets) for 8 weeks; control mice were of the same genotype. Durations of diet and antibody therapies varied as outlined in the main text.

##### Model of streptozotocin-induced diabetes and advanced NASH

We engaged contract research organization (CRO) service from SMC Laboratories, Japan to perform this study. Briefly, two-day old male wild-type mice received a single subcutaneous injection of 200 µg streptozotocin (STZ, S1030, Sigma), followed by feeding with high fat diet (HFD32, CLEA Japan) from when they were of 4-weeks of age until the end of the experiment at 18-weeks. Control mice received NC diet for the duration of the experiment. Mice received either IgG, X203 or X209 from week 11 until the end of the experiment.

##### II11ra-deleted mice

Mice lacking functional alleles for Il11rα (*II11ra^-/-^*) were on C57Bl/6J genetic background (B6.129S1-*II11ra^tm1Wehi^*/J, Jackson’s Laboratory). Both *Il11ra^-/-^* mice and their wild-type littermates (*II11ra^+/+^*) were fed with HFMCD for 10 weeks from 5-weeks of age to develop NASH; control mice were fed with NC for the same duration.

### *In vivo* administration of Il-11

Recombinant mouse Il-11 (rmil-11) was reconstituted to a concentration of 50 µg ml^-1^ in saline. Ten-week-old male *Col1a1-GFP* reporter mice^1^ and wild-type C57BL/6J mice received daily subcutaneous injection of either 100 µg kg^-1^ of rmIl-11 or identical volume of saline for 21 days.

### *In vivo* administration of anti-IL-11 or anti-IL11RA monoclonal antibodies

Mice were injected intraperitoneally with either anti-IL-11 (X203) or anti-IL11RA (X209) or an identical amount of IgG isotype control for the treatment durations outlined in the main text.

### Antibodies

ACTA2 (ab7817, Abcam), CD45 (103102, Biolegend), Collagen I (ab34710, Abcam), ERK1/2 (4370, Cell Signaling), ERK1/2 (4695, Cell Signaling), GAPDH (2118, Cell Signaling), IgG (Aldevron), IL-11 (X203, Aldevron), IL11RA (X209, Aldevron), Ly6C (128039, Biolegend), TGFβ1 (141402, Biolegend), anti rabbit HRP (7074, CST), anti mouse HRP (7076, CST).

### Recombinant proteins

Commercial recombinant proteins: Human angiotensin II (A9525, Sigma-Aldrich), human CCL2 (279-MC-050/CF, R&D Systems), human bFGF (233-FB-025, R&D Systems), human IL-11 (PHC0115, Life Technologies), human PDGF (220-BB-010, R&D Systems), human TGFβ1 (PHP143B, Bio-Rad), and mouse TGFβ1 (7666-MB-005, R&D Systems).

Custom recombinant proteins: Mouse Il-11 (UniProtKB: P47873) were synthesized without the signal peptide, HyperIL-11 was constructed using a fragment of IL11RA (amino acid residues 1–317; UniProtKB: Q14626) and IL-11 (amino acid residues 22–199, UniProtKB: P20809) as described previously^3^. All custom recombinant proteins were synthesized by GenScript using a mammalian expression system.

### Chemical

Hydrogen Peroxide (H_2_O_2_, 31642, Sigma)

### Cell culture

Cells (HSCs or fibroblasts) were grown and maintained at 37°C and 5% CO_2_. The growth medium was renewed every 2–3 days and cells were passaged at 90% confluence using standard trypsinization techniques. All the experiments were carried out at low cell passage (<P4) and cells were serum-starved for 16 h prior to respective stimulations. Stimulated cells were compared to unstimulated cells that have been grown for the same duration under the same conditions (serum-free media), but without the stimuli.

#### Primary human atrial fibroblasts

Human atrial fibroblasts were prepared and cultured as described previously^2^.

#### Primary human hepatic stellate cells (HSCs)

HSCs (5300, ScienCell) were cultured in stellate cells complete media (5301, ScienCell) on poly-L-lysine-coated culture plates (2 µg cm^-2^, 0403, ScienCell).

### Operetta high throughput phenotyping assay

The Operetta phenotyping assay was performed mostly as described previously^3^ with minor modifications described here: HSCs were seeded in 96-well black CellCarrier plates (PerkinElmer) at a density of 5×10^3^ cells per well. Following experimental conditions, cells were fixed in 4% paraformaldehyde (PFA, 28908, Thermo Fisher Scientific), permeabilized with 0.1% Triton X-100 (Sigma) and non-specific sites were blocked with 0.5% BSA and 0.1% Tween -20 in PBS. Cells were incubated overnight (4°C) with primary antibodies (1:500), followed by incubation with the appropriate AlexaFluor 488 secondary antibodies (1:1000). EdU-Alexa Fluor 488 was incorporated using a Click-iT EdU Labelling kit (C10350, Thermo Fisher Scientific) according to manufacturer’s protocol. Cells were counterstained with 1 µg ml^-1^ DAPI (D1306, Thermo Fisher Scientific) in blocking solution. Each condition was imaged from duplicated wells and a minimum of 7 fields per well using Operetta high-content imaging system 1483 (PerkinElmer). The quantification of ACTA2^+ve^ cells was measured using Harmony v3.5.2 (PerkinElmer). The measurement of fluorescence intensity per area of Collagen I (normalized to the number of cells) was performed with Columbus 2.7.1 (PerkinElmer).

### Matrigel invasion assay

The invasive behavior of human HSCs was assayed using 24-well Boyden chamber invasion assays (Cell Biolabs Inc.). Equal numbers of HSCs in serum-free HSC media were seeded in triplicates onto the ECM-coated matrigel and were allowed to invade towards HSC media containing 0.2% FBS. After 48 h of incubation with stimuli, media was aspirated and non-invasive cells were removed using cotton swabs. The cells that invaded towards the bottom chamber were stained with cell staining solution (Cell Biolabs Inc.) and invasive cells from 5 non-overlapping fields of each membrane were imaged and counted under 40x magnification. For antibody inhibition experiments, HSCs were pretreated with X203, X209, or IgG control antibodies for 15 m prior to addition of stimuli.

### Generation of mouse monoclonal antibodies against IL11RA

#### Genetic immunisation and screening for specific binding

A cDNA encoding amino acids 23-422 of human IL11RA was cloned into expression plasmids (Aldevron). Mice were immunised by intradermal application of DNA-coated gold-particles using a hand-held device for particle-bombardment. Cell surface expression on transiently transfected HEK cells was confirmed with anti-tag antibodies recognising a tag added to the N-terminus of the IL11RA protein. Sera were collected after 24 days and a series of immunisations and tested in flow cytometry on HEK293 cells transiently transfected with the aforementioned expression plasmids. The secondary antibody was goat anti-mouse IgG R-phycoerythrin-conjugated antibody (Southern Biotech, #1030-09) at a final concentration of 10 µg ml^-1^. Sera were diluted in PBS containing 3% FBS. Interaction of the serum was compared to HEK293 cells transfected with an irrelevant cDNA. Specific reactivity was confirmed in 2 animals and antibody-producing cells were isolated from these animals and fused with mouse myeloma cells (Ag8) according to standard procedures. Supernatant of hybridoma cultures were incubated with HEK cells expressing an IL11RA-flag construct and hybridomas producing antibodies specific for IL11RA were identified by flow cytometry.

#### Identification of neutralizing anti-IL11RA antibodies

Antibodies that bound to IL11RA-flag cells but not to the negative control were considered specific binders and subsequently tested for anti-fibrotic activity on human and mice atrial fibroblasts as described by Schafer et al^2^. Briefly, primary human or mouse fibroblasts were stimulated with human or mouse TGFβ1, respectively (5 ng ml^-1^; 24 h) in the presence of the antibody candidates (6 µg ml^-1^). TGFβ1 stimulation results in an upregulation of endogenous IL-11, which if neutralized, blocks the pro-fibrotic effect of TGFβ1. The fraction of activated myofibroblasts (ACTA2^+ve^ cells) was measured on the Operetta platform as described above to estimate the neutralization potential of the antibody candidates. In order to block potential trans-signalling effects, antibodies were also screened in the context of hyperIL-11 stimulation of human fibroblasts (200 pg ml^-1^). We detected three specific and neutralizing IL11RA antibodies, of which X209 was taken forward for *in vivo* studies. The same procedures were performed to obtain a neutralizing antibody that binds to the ligand IL-11, as detailed by Cook *et al*^3^.

#### Binding kinetics of X209 to IL11RA

Binding of X209 to human IL11RA was measured on Biacore T200 (GE Healthcare). X209 was immobilized onto an anti-mouse capture chip. Interaction assays were performed with HEPES-buffered saline pH 7.4 containing 0.005% P20 and 0.5% BSA. A concentration range (1.56 nM to 100 nM) of the analyte (human IL11RA) was injected over X209 and reference surfaces at a flow rate of 40 μl min^-1^. Binding to mouse Il11ra1 was confirmed on Octet system (ForteBio) using a similar strategy. All sensograms were aligned and double-referenced^5^. Affinity and kinetic constants were determined by fitting the corrected sensograms with 1:1 Langmuir model. The equilibrium binding constant *K*_D_ was determined by the ratio of *k*_d_/*k*_a_.

#### X209 IC_50_ measurement

HSCs were stimulated with TGFβ1(5 ng ml^-1^, 24 h) in the presence of IgG (4 µg ml^-1^) and varying concentrations of X209 (4 µg ml^-1^ to 61 pg ml^-1^; 4-fold dilutions). Supernatants were collected and assayed for the amount of secreted MMP2. Dose-response curves were generated by plotting the logarithm of X209 tested concentration (pM) versus corresponding percent inhibition values using least squares (ordinary) fit. The IC_50_ value was calculated using log(inhibitor) versus normalized response-variable slope equation.

#### Blood pharmacokinetics and biodistribution

C57BL/6J mice (10-12-weeks old) were retro-orbitally injected (left eye) with 100 µl of freshly radiolabeled ^125^I-X209 (5µCi, 2.5 µg) in PBS. Mice were anesthetized with 2% isoflurane and blood were collected at several time points (2, 5, 10, 15, 30 m, 1, 2, 4, 6, 8 h, 1, 2, 3, 7, 14 and 21 days) post injection via submandibular bleeding. For biodistribution studies, blood was collected via cardiac puncture and tissues were harvested at the following time points: 1, 4 h, 1, 3, 7, 14, 21 days post injection. The radioactivity contents were measured using a gamma counter (2480 Wizard2, Perkin Elmer) with decay-corrections (100x dilution of 100 μl dose). The measured radioactivity was normalized to % injected dose/g tissue.

### Precision cut liver slices (PCLS) and Western blotting of NASH patient liver

We engaged CRO service (FibroFind, UK) to perform these studies. Briefly, human PCLS were cut and incubated with TGFβ1 for 24 h. ELISA from the supernatant was performed using Human IL-11 DuoSet (DY218, R&D Systems). This CRO also collected liver biopsies from patients undergoing liver resections for cancers where adjacent, non-cancerous tissue was collected for molecular studies. Patients had either no documented intrinsic liver disease (controls) or previously documented alcoholic liver disease, primary biliary cirrhosis, primary sclerosing cholangitis or NASH. For confidentiality reasons no further information was provided for these samples.

### RNA-seq

#### Generation of RNA-seq libraries

Total RNA was quantified using Qubit RNA high sensitivity assay kit (Thermo Fisher Scientific) and RNA integrity number (RIN) was assessed using the LabChip GX RNA Assay Reagent Kit (Perkin Elmer). TruSeq Stranded mRNA Library Preparation Kit (Illumina) was used to prepare the transcript library according to the manufacturer’s protocol. All final libraries were quantified using KAPA library quantification kits (KAPA Biosystems). The quality and average fragment size of the final libraries were determined using LabChip GX DNA High Sensitivity Reagent Kit (Perkin Elmer). Libraries were pooled and sequenced on a NextSeq 500 benchtop sequencer (Illumina) using NextSeq 500 High Output v2 kit and paired-end 75-bp sequencing chemistry.

#### RNA-seq analysis

##### Stiffness-induced RNA regulation in hepatic stellate cells

Normalized gene expression values were downloaded from Dou et al^5^. Lowly expressed genes (FPKM at baseline ≥ 2) were removed from the analysis and fold changes were calculated as average FPKM in HSCs on stiff surface divided by average FPKM in HSCs on soft surface. The fold change of RNA expression for upregulated genes (f.c. >1) was plotted and genes were ranked according to their average FPKM value. *TGFB1 stimulation of human hepatic stellate cells and antibody treatment in HFMCD:* Sequenced libraries were demultiplexed using bcl2fastq v2.19.0.316 with the *—no-lane-splitting* option. Adapter sequences were then trimmed using trimmomatic^6^ v0.36 in paired end mode with the options *MAXINFO:35:0.5 MINLEN:35*. Trimmed reads were aligned to the *Homo sapiens* GRCh38 using STAR^7^ v. 2.2.1 with the options *—outFilterType BySJout—outFilterMultimapNmax 20 — alignSJoverhangMin 8 — alignSJDBoverhangMin 1 — outFilterMismatchNmax 999 — alignIntronMin 20 — alignIntronMax 1000000 — alignMatesGapMax 1000000* in paired end, single pass mode. Only unique alignments were retained for counting. Counts were calculated at the gene level using the FeatureCounts module from subread^8^ v. 1.5.1, with the options *-O -s 2 -J -T 8 -p -R -G*. The Ensembl release 92 hg38 GTF was used as annotation to prepare STAR indexes and for FeatureCounts. For the antibody treatment experiments in mouse, libraries were treated as for the human samples, only using mm10 Ensembl release 86 genome and annotation. Differential expression analyses were performed in R 3.4.1 using the Bioconductor package DESeq29^9^ 1.18.1, using the Wald test for comparisons and including the variance shrinkage step setting a significance threshold of 0.05. Gene set enrichment analyses (GSEA) were performed in R 3.4.1 using the fgsea package and the MSigDB Hallmark genesets^10,11^, performing 100000 iterations. The “stat” column of the DESeq2 results output was used as ranked input for each enrichment, taking only mouse genes with one-to-one human orthologs.

### Mass cytometry by Time of Flight (CyTOF)

Immune cells were isolated from liver as described previously^12^. Liver tissues were minced and digested with 100 µg ml^-1^ Collagenase IV and 20 U ml^-1^ DNase I, at 37°C for 1 h. Following digestion, cells were passed through strainer to obtain single cell suspension and subjected to percoll gradient centrifugation for isolation of immune cells. CyTOF staining was performed as previously described^13^. Cells were thawed and stained with cisplatin (Fluidigm) to identify live cells, followed by staining with metal-conjugated CD45 antibody, for barcoding purpose. After barcoding, cells were stained with metal-conjugated cell surface antibody (Ly6C). Cells were then fixed with 1.6% PFA, permeabilized with 100% methanol, and subjected to intracellular antibody staining (TGFβ1). Cells were finally labeled with DNA intercalator before acquisition on Helios mass cytometer (Fluidigm). For analysis, first live single cells were identified, followed by debarcoding to identify individual samples. Manual gating was performed using Flowjo software (Flowjo, LLC, USA).

### Enzyme-linked immunosorbent assay (ELISA) and colorimetric assays

The levels of IL-11 and MMP-2 in equal volumes of cell culture media were quantified using Human IL-11 Quantikine ELISA kit (D1100, R&D Systems) and Total MMP-2 Quantikine ELISA kit (MMP200, R&D Systems), respectively. Mouse serum levels of alanine aminotransferase (ALT) was measured using Alanine Transaminase Activity Assay Kit (ab105134, abcam). Total secreted collagen in the cell culture supernatant was quantified using Sirius red collagen detection kit (9062, Chondrex). Total hydroxyproline content in the livers was measured using Quickzyme Total Collagen assay kit (Quickzyme Biosciences). Liver Triglycerides (TG) measurements were performed using triglyceride colorimetric assay kit (10010303, Cayman). All ELISA and colorimetric assays were performed according to the manufacturer’s protocol.

### Quantitative polymerase chain reaction (qPCR)

Total RNA was extracted from either the snap-frozen liver tissues or HSCs lysate using Trizol (Invitrogen) followed by RNeasy column (Qiagen) purification. The cDNAs were synthesized with iScript^TM^ cDNA synthesis kit (Bio-Rad) according to manufacturer’s instructions. Gene expression analysis was performed on duplicate samples with either TaqMan (Applied Biosystems) or fast SYBR green (Qiagen) technology using StepOnePlus^TM^ (Applied Biosystem) over 40 cycles. Expression data were normalized to *GAPDH* mRNA expression and fold change was calculated using 2^-∆∆Ct^ method. The sequences of specific TaqMan probes and SYBR green primers are available upon request.

### Immunoblotting

Western blots were carried out on total protein extracts from HSCs and liver tissues. Both cells and frozen tissues were homogenized in radioimmunoprecipitation assay (RIPA) buffer containing protease and phosphatase inhibitors (Thermo Scientifics), followed by centrifugation to clear the lysate. Protein concentrations were determined by Bradford assay (Bio-Rad). Equal amount of protein lysates were separated by SDS-PAGE, transferred to PVDF membrane, and subjected to immunoblot analysis for the indicated primary antibodies. Proteins were visualized using the ECL detection system (Pierce) with the appropriate secondary antibodies.

### Histology

Liver tissues were fixed for 48 h at RT in 10% neutral-buffered formalin (NBF), dehydrated, embedded in paraffin blocks and sectioned at 7μm. Sections stained with Masson’s Trichrome were examined by light microscopy.

### Statistical analysis

Statistical analyses were performed using GraphPad Prism software (version 6.07). Fluorescence intensity (Collagen I) was normalized to the number of cells detected in the field and recorded for 7 fields per well. Cells expressing ACTA2 were quantified and the percentage of activated fibroblasts (ACTA2^+ve^) was determined for each field. P values were corrected for multiple testing according to Dunnett’s (when several experimental groups were compared to one condition), Tukey (when several conditions were compared to each other within one experiment), Sidak (when several conditions from 2 different genotypes were compared to each other). Analysis for two parameters (antibody efficacy across time) for comparison of two different groups were performed by two-way ANOVA. The criterion for statistical significance was *P* <0.05.

### Data Availability

High-throughput sequencing data generated for this study can be downloaded from the (GEO) repository (data currently under submission). All other data are in the manuscript or in the supplementary materials.

